# Chemical control of 2’-hydroxyl-dependent Cas9 target engagement enables CRISPR RNA ribose replacement

**DOI:** 10.64898/2026.01.26.701763

**Authors:** Adrian A. Pater, Halle M. Barber, Sruthi Sudhakar, Ramadevi Chilamkurthy, Sunit K. Jana, Mansi A. Parasrampuria, Michael. S. Bosmeny, Jacob A. Graczyk-Marrs, Seth B. Eddington, Cole A. Blazier, Leonora Abdullahu, Elise Malek-Adamian, Christopher L. Barkau, Daniel O’Reilly, Sergey Korolev, P.I. Pradeepkumar, Masad J. Damha, Keith T. Gagnon

**Affiliations:** Department of Biochemistry, Wake Forest University School of Medicine, Winston-Salem, NC, USA; Department of Chemistry, McGill University, Montreal, Quebec, Canada; Department of Chemistry, Indian Institute of Technology Bombay, Powai, Mumbai, India; Chemistry & Biochemistry, Southern Illinois University, Carbondale, IL, USA; Biochemistry & Molecular Biology, Southern Illinois University, School of Medicine, Carbondale, IL, USA; Biochemistry & Molecular Biology, Saint Louis University, Saint Louis, MO, USA

**Keywords:** Cas9, guide RNA, CRISPR, chemical modification, ribose, 2’-hydroxyl, checkpoint, target engagement, therapeutics, gene editing, gene therapy, medicine, genome engineering

## Abstract

Advanced CRISPR-based therapies benefit from CRISPR RNA (crRNA) with high nuclease resistance and enhanced drug-like properties, which is primarily achieved through chemical replacement of the RNA ribose moiety. However, for gene editing enzymes like CRISPR-Cas9 a handful of residues cannot be replaced with chemical ribose analogues, limiting the scope of therapeutic strategies. The mechanism underlying this restriction has remained unclear. Here, using nucleic acid chemistry, biochemistry, cryo-EM, and molecular dynamics simulations, we show that the ribose 2’-hydroxyl group at specific crRNA residues is required to achieve a conformational state competent for Cas9 target DNA binding. Based on the mechanistic principles uncovered, we combined site-specific phosphorothioate linkage chemistry with ribose replacement chemistry to restore binding and activity, resulting in high Cas9 editing efficiency and fidelity with a ribose-free crRNA. This study offers novel mechanistic insight and crRNAs with full chemical stabilization, making rational design of guide RNAs with complete nuclease protection for CRISPR-based medicines possible.

## Introduction

Clustered regularly interspaced short palindromic repeats (CRISPR)-associated (Cas) endonucleases and their cognate CRISPR RNA (crRNA) guides are being harnessed to potentially treat a variety of genetic diseases through sequence-specific gene editing. *Streptococcus pyogenes* Cas9 (SpCas9) is one of the most well-characterized CRISPR-Cas systems being developed for human therapeutics ^1, 2^. The naturally occurring CRISPR-Cas9 complex comprises a Cas9 endonuclease and a guide RNA, composed of a duplex between two RNAs, the crRNA and *trans*-activating crRNA (tracrRNA). Alternatively, the two guides are commonly fused into a single guide RNA (sgRNA) ^3^. Together, these components assemble into a ribonucleoprotein (RNP) complex. The assembled RNP is capable of binding and cleaving target DNA that is complementary to the guide RNA spacer sequence. These biochemical mechanisms are the basis for developing CRISPR-based therapeutics, ranging from gene knockout with traditional CRISPR-Cas9 to more precise technologies that fuse Cas9 to diverse enzymes, such as base editing ^4–7^ and prime editing ^5, 8, 9^.

For therapeutic applications, CRISPR-Cas can be delivered through viral or non-viral methods ^10–12^. Viral delivery approaches like adeno-associated viruses (AAV) offer some advantages, such as high efficiency and robust expression ^13, 14^. However, disadvantages include limited cargo capacity ^10^, risks from genome integration ^15–17^, and long-term expression of CRISPR-Cas components, which can induce off-target editing or elimination of edited cells by the immune system ^15, 18, 19^. A leading, transient non-viral alternative is to encapsulate Cas9-encoding mRNA and guide RNA in lipid nanoparticles (LNP) ^11, 20, 21^. While attractive, LNPs are not entirely effective at preventing immunogenicity^22^ or protecting RNA from intracellular degradation ^23^. LNPs also remain difficult to engineer with high tissue or cell specificity ^24^ and require robust yet safe release from endosomes, a process that is still not fully understood ^25^.

For nucleic acid therapeutics that have achieved regulatory approval in humans, complete or near complete chemical modification at every nucleotide position has proven essential to achieve high marks for safety and potency^26–29^. The latest generation of small interfering RNA (siRNA) drugs are completely chemically stabilized, ribose-free, and possess tissue-targeting conjugates like *N*-acetyl galactosamine (GalNAc) ^30^. These provide safe and low doses, delivered subcutaneously as infrequently as once every 6 months ^31^, and no longer rely on LNPs for delivery. Although CRISPR-Cas systems require both a protein component and guide RNA, separate delivery of the guide RNA, similar to current state-of-the-art siRNAs, could offer new forms of hybrid delivery. These might include genetic encoding of Cas9 and tracrRNA by viral vectors, followed by separate delivery of a fully chemically protected crRNA with a tissue-targeting conjugate ^32, 33^. Advantages might include better spatiotemporal control of editing, delivery to difficult tissues or cells, sequential editing at multiple genes, in multiple tissues, or at different times, safer redosing schemes, and subsequent turn-off or removal of CRISPR or AAV components once editing is complete ^33^. However, exploring such alternative delivery modalities has not been possible due to rapid degradation of guide RNAs *in vivo* and the inability to fully chemically stabilize them ^26, 34, 35^.

Modifications like 2’-*O*-methyl (2’-OMe) with phosphorothioate (PS) linkages placed at guide RNA termini have improved exonuclease stability and internal modifications ^36, 37^, including DNA (2’-H) and bridged nucleic acids (BNAs), have enhanced specificity ^38–41^. However, these modifications are insufficient to protect against endonucleases and attempts to fully chemically modify crRNA significantly reduce Cas9 activity ^33, 38, 42–45^. Structural inspection revealed that the most sensitive nucleotides engage in hydrogen bonding between their 2’-hydroxyl (2’-OH) group and the Cas9 peptide backbone ^43, 46^. Thus, the challenge in fully replacing all ribose moieties in guide RNAs has been referred to as the “2’-hydroxyl barrier” ^47^. A similar barrier has been reported for CRISPR-Cas12a ^48, 49^. Understanding why these interactions are important might reveal new mechanistic insight for CRISPR-Cas regulation and enable full ribose replacement and chemical tuning of crRNAs.

Several strategies have been explored to overcome the 2’-hydroxyl barrier. One was the placement of PS linkages 3’ of sensitive 2’-ribose positions, rather than ribose replacement, to protect from nucleases ^50^. However, PS modifications have been shown to compromise activity when flanking 2’-OH contact residues ^42^, do not fully stabilize against nucleases ^51^, and may be toxic if overused ^52^. A recent approach utilized a fully chemically modified protecting oligo complementary to and overlapping with critical 2’-OH contact residues ^53^. While promising, this method did not recapitulate full editing efficiency *in vivo*. Thus, finding a viable modification strategy that maintains high editing efficiency while conferring desired drug-like properties will likely require a deeper mechanistic understanding of the principles underlying the 2’-hydroxyl barrier.

Here we sought to determine the influence of ribose replacement chemistries on Cas9-mediated DNA editing, target binding, RNP stability, and conformational states with an emphasis on the role of the 2’-OH at critical contact residues. We systematically probed structure-function relationships at 2’-OH positions using various sugar modifications and examined their effects on Cas9 biochemical properties and gene editing. We found that 2’-OH contacts allosterically mediate a Cas9 conformational state that is competent for DNA target binding. Based on these principles, we rescued activity of a ribose-free crRNA by careful placement of PS linkages adjacent to specific 2’-OH contact positions, which we hypothesized would modulate interaction dynamics with Cas9. This resulted in a ribose-free crRNA design, FM8.15, that shows favorable target binding, editing efficiency, and specificity. These results also implicate 2’-OH contacts as critical mediators of a previously unknown target engagement checkpoint mechanism for Cas9.

## Materials and Methods

### Synthesis of guide RNAs

Standard phosphoramidite solid-phase synthesis conditions were employed to synthesize both modified and unmodified oligonucleotides. The syntheses were carried out on an Applied Biosystems 3400, or an Expedite DNA Synthesizer at a scale of 1 micromole utilizing Unylink CPG support (ChemGenes). All phosphoramidites were prepared as 0.15 M solutions in acetonitrile (ACN), with the exception of DNA, which was prepared as 0.1 M solutions. A 0.25 M solution of 5-ethylthiotetrazole in ACN was used to activate the phosphoramidites for the coupling step. Detritylation was conducted using 3% trichloroacetic acid in CH_2_Cl_2_ for 110 s. Capping incomplete sequences was performed with acetic anhydride in tetrahydrofuran (THF) alongside 16% N-methylimidazole in THF. Oxidation was achieved using 0.1M I_2_ in the ratio of 1:2:10 pyridine, water, and THF. The coupling durations for phosphoramidites ranged from 10 to 15 min. Deprotection and cleavage from the solid support were performed with either a 3:1 mixture of NH_4_OH and EtOH for 48 h at room temperature (RT), or at 55°C for 16 h. Oligonucleotides containing RNA were synthesized using standard 2’-TBDMS phosphoramidites; desilylation was accomplished with either neat triethylamine trihydrofluoride for 48 h at RT, or with a mixture of triethylamine trihydrofluoride, N-methyl pyrrolidone, and triethylamine in a volume ratio of 1.5:0.75:1 for 2.5 h at 65°C. Crude oligonucleotides were purified with a Waters 1525 anion-exchange HPLC using a Source 15Q resin column (11.5 cm x 3 cm), flow rate of 10 mL/min, and a gradient of 0-100% buffer B (0.5M LiClO_4_) over 50 min at 60°C. Samples were desalted using NAP-25 desalting columns following the manufacturer’s instructions. All crRNAs contained a 19 nt spacer and oligonucleotides used in this study, excluding primers, are listed in **Table S1**. Modified crRNAs were prepared for RNP assembly by heating them to 95°C and then placing them on ice to inhibit the formation of stable secondary structures.

### Cas9 protein expression and purification

pET-Cas9-NLS-6xHis (Addgene #62933) plasmid encoding SpCas9, or a catalytically inactive “dead” Cas9 (dCas9) plasmidprepared by site-directed mutagenesis to generate H840A and D10A mutations (pET-dCas9-NLS-6xHis), were used. Protein expression was induced with 0.4 mM isopropyl β-D-1-thiogalactopyranoside (IPTG) and expressed for 16 h at 18°C in *Escherichia coli* Rosetta 2 (DE3) (Novagen, Cat. #71397-3) cells. Cells were resuspended in 12 mL of chilled binding buffer (20 mM Tris–HCl, pH 7.5, 250 mM NaCl, 1 mM PMSF, 5 mM imidazole) per 500 mL of culture pellet and lysed via French press at 1,000-1,100 psi repeated twice. Lysate was clarified by centrifugation, loaded on a HisPur™ Cobalt Resin (ThermoFisher Scientific, Cat. #71397-3) column, and washed sequentially with 20 mM Tris–HCl, pH 7.5, containing increasing NaCl concentrations (0.25/0.5/0.75/1.0 M) and 20 mM imidazole. Protein was eluted with 15 mL elution buffer (20 mM Tris-HCl, pH 7.5, 250 mM NaCl, and 130 mM imidazole) and concentrated and buffer exchanged (20 mM Tris-HCl, pH 7.5, 150 mM KCl, 1 mM EDTA, 1 mM DTT) using Vivaspin 15 centrifugal concentrators (Sartorius, 15K MWCO). The concentrated protein was applied to a Superdex 200 10/300 GL column (GE Healthcare) and eluted with 20 mM Tris-HCl, pH 7.5, 150 mM KCl, 1 mM EDTA, 1 mM DTT. “Fraction 13” containing Cas9 of the apparent correct size and free of nucleic acid contamination was collected, concentrated, and buffer exchanged into 40 mM Tris-HCl, pH 7.5, 300 mM KCl, 2 mM EDTA, 2 mM DTT using Vivaspin 15 centrifugal concentrators. One volume of glycerol was added for a final 50% (v/v) concentration. Protein concentration was determined using a UV spectrophotometer at 280 nm with the extinction coefficient 120,450 M^-1^ cm^-1^ and calculated using Beer–Lambert law.^54^

### *In vitro* Cas9 cleavage activity assays

*In vitro* cleavage assays were performed as previously described ^38, 44^ with minor modifications. Cleavage reactions were carried out at 37°C in 1x cleavage buffer (20 mM Tris-HCl, pH 7.5, 100 mM KCl, 5% glycerol, 1 mM DTT, 0.5 mM EDTA, 2 mM MgCl_2_) using 160 ng of EGFP PCR-amplified DNA, 0.1 mg/mL of purified yeast tRNA, and Cas9 ribonucleoprotein (RNP) complex containing 0.25 µM SpCas9, 0.3 µM crRNA, and 0.25 µM tracrRNA. Reactions were incubated for 2 h at 37°C to ensure complete cleavage. Reactions were treated sequentially with 10 µg RNase A (10 min) and 10 µg Proteinase K (10 min) to degrade residual RNA and proteins. DNA was precipitated by adding 10 volumes of 2% LiClO_4_ in 100% acetone and incubated overnight at -20°C. The precipitated sample was resolved by electrophoresis on a 1.5% agarose gel stained with ethidium bromide. Gels were visualized using the ChemiDoc MP Imaging System (BioRad), and cleavage bands were quantified using ImageJ to determine the fraction cleaved.^55^ Time course cleavage assays were performed in the same way, however reactions were terminated with the addition of 2% LiClO_4_ in 100% acetone at indicated time points, precipitated at -20°C, resuspended, and treated sequentially with RNase A (10 min) and Proteinase K (10 min) as described above. Observed rate constants were calculated by fitting time course cleavage data by non-linear regression in GraphPad Prism to an exponential one-phase association (crEG-SJ3 and crEG-SJ16) or two-phase association (crEG, crEG-SJ1, and crEG-SJara) equation. For one-phase association, an observed rate (*k*_obs_) is provided directly from calculations. For two-phase association, an average *k*_obs_ is estimated from the equation *k*_obs_ = [((*k*_fast_)*(%Fast))+((*k*_slow_)*(%Slow))]/100, which can be considered as the mean reaction time ^56^.

### Cell editing quantified by EGFP knockout and flow cytometry

HEK293T cells stably expressing EGFP and SpCas9 were kindly provided by the Dr. Wen Xue (UMass Chan Medical School) ^57^. Cell-based gene editing of Cas9 was carried out as previously described ^44^. Briefly, cells were cultured in 1x minimum essential medium (MEM, Corning 10-010-CV) supplemented with 5% fetal bovine serum (FBS, Gibco), 5% cosmic calf serum (CCS, Gibco), and 1% non-essential amino acids (NEAA, Gibco). Cells were maintained at 37°C and 5% CO_2_. For transfections, 40,000 cells were reverse-transfected with 5 pmol of 1:1 molar ratio of crRNA:tracrRNA using 0.5 µL of Lipofectamine RNAiMAX (ThermoFisher Scientific) diluted in Opti-MEM (Gibco) to a final volume of 200 µL. Lipid complexes were incubated for 20 min following the manufacturer’s recommended protocol then added to cells. After 8 h of incubation, one volume of full serum media (5% FBS, 5% CCS and 1% NEAA) was added and cells were incubated for an additional 16 h. After this period, the medium was replaced with full serum medium, and the transfected cells were grown for an additional 4 d with media change at 2 d.

For experiments testing Cas9 Y450A and F105A, mutants were generated by site-directed mutagenesis of LentiV_Cas9_Puro (Addgene #108100). For plasmid electroporation, 2 × 10^6^ HEK293T cells stably expressing EGFP were electroporated with 8 µg of plasmid in a T75 flask using the Neon NxT electroporation system (ThermoFisher Scientific) and the 100 μL Neon Transfection System Kit (ThermoFisher Scientific, Cat. #MPK10096) per manufacturer’s HEK293 protocol (1,100 V, 20 ms per pulse, 2 pulses). After 36 h of recovery time, 50,000 cells were reverse transfected with 10 pmol of crRNA:tracrRNA duplex in Lipofectamine RNAiMAX (ThermoFisher Scientific, Cat. #13778100), diluted in Opti-MEM (ThermoFisher Scientific, Cat. #11058021) to a final volume of 200 µL, and incubated for 20 min as per the manufacturer’s recommended protocol. The lipid complex was added to each well and incubated for 12 h. Opti-MEM was replaced with full serum media and cells were cultured for an additional 4.5 d with media change at 2 d.

At 5 d post-transfection, cells were harvested for flow cytometry analysis. Cells were washed with 1x PBS (Gibco), trypsinized, and centrifuged at 500 x g for 5 min. The supernatant was removed, and the cell pellet was washed with 1x PBS. The cells were then centrifuged again, and the supernatant was removed. The cell pellet was resuspended in 1x PBS and transferred to a round-bottom 96-well plate. Resuspended cells were analyzed using an Attune NxT Flow Cytometer (ThermoFisher Scientific) or LSRFortessa X-20 (BD Biosciences). Forward and side scatter were applied for doublet discrimination. EGFP fluorescence was measured using a 488 nm laser for EGFP excitation. Data acquisition was acquired on FACSDiva (BD FACSDiva), and analysis was performed on FCS Express (Dotmatics).

### Thermal denaturation of RNPs

Thermal denaturation experiments were performed using a Cary 3500 Multicell UV-vis spectrophotometer (Agilent). Reaction mixtures containing Cas9 alone and Cas9, tracrRNA and modified crRNAs were prepared in 1x UV-melt buffer (20 mM Cacodylate, pH 7.5, 150 mM KCl, 1 mM MgCl_2_) at a final equimolar concentration of 1 µM. The reactions were incubated at RT for 10 min. The incubated samples were transferred to a quartz cuvette (Starna, Cat. #26.100/LHS/Q/10) and UV-absorbance measured at 280 nm every 0.2°C, over the range of 32-68°C with a ramp rate of 1°C/min. The melting temperature (*T*_m_) was determined using the onboard Cary UV Workstation (v.1.2.328) software via the first derivative method.

### DNA target binding measured by dot blot filter binding assays

Radiolabeling of target DNA for dot blots was performed as previously described ^58^ with minor modifications. Briefly, 100 pmol of antisense DNA target strand was radiolabeled with γ-^32^P-ATP using T4 polynucleotide kinase (ThermoFisher Scientific) according to the manufacturer’s protocol. Labeled oligonucleotides were purified by phenol-chloroform extraction and ethanol precipitation, then annealed to sense DNA strand at 1.2-fold molar excess at 95°C for 3 min, followed by slow cooling. The duplex was resolved on a 15% non-denaturing polyacrylamide gel and purified using the crush-and-soak gel purification method. Radioactivity of purified duplex was quantified by scintillation counting.

To assess dCas9 binding, a 1:1.2:1.2 molar ratio of Cas9:tracrRNA:crRNA was assembled in 1x cleavage buffer (20 mM Tris-HCl, pH 7.5, 100 mM KCl, 5% glycerol, 1 mM DTT, 0.5 mM EDTA, 2 mM MgCl_2_, 0.1 mg/mL of purified yeast tRNA, 0.5 mg/mL of BSA) to form the ribonucleoprotein (RNP) complex. The γ-^32^P-labeled duplex DNA (1000 cpm per reaction) was incubated with increasing concentrations of the RNP in 1x cleavage buffer at 37°C for 15 min. Reactions were gently vacuum-filtered through a nitrocellulose membrane using a 96-well dot blot apparatus (GE Whatman), washed twice with 1x cleavage buffer, and washed 3x with 1x PBS (Corning). Membranes were visualized using a Typhoon FLA 9500 phosphorimager and radioactive spots were quantified with ImageQuant software and plotted in GraphPad Prism (v.10.4.2). Data was fit to a two-site binding model for visualization, but *K*_d_ and *B*_max_ values were estimated using a one-site binding model.

### DNA target binding measured by CRISPRa

The vector pMA2780 lenti tet on EGFP (Addgene #128043) was packaged into lentiviral particles by VectorBuilder. HeLa Tet-On 3G cells (Takara, Cat. #631183) were transduced according to the manufacturer’s recommended protocol. After 5 days, EGFP expression was induced with 0.5 µg/mL doxycycline and stable cells selected by fluorescence activated cell sorting (FACS) based on EGFP fluorescence.

For CRISPRa transfections, HeLa Tet-On 3G EGFP cells were cultured in 1x MEM supplemented with 10% tetracycline-free FBS and 1% NEAA. Cells were maintained at 37°C and 5% CO_2_. To introduce dCas9-VPR, 2 x 10^6^ cells were electroporated with 8 μg of dCas9-VPR_P2A_mCherry (Addgene #154193) using the Neon NxT electroporation system (ThermoFisher Scientific) and the 100 μL Neon Transfection System Kit (ThermoFisher Scientific, Cat. #MPK10096) using a program of 1,005 V, 35 ms per pulse, 2 pulses. After 48 h, 75,000 cells were reverse transfected with 20 pmol of 1:1 molar ratio of crRNA:tracrRNA duplex using 1 μL Lipofectamine RNAiMAX (ThermoFisher Scientific, Cat. #13778100), diluted in 400 μL Opti-MEM (ThermoFisher Scientific, Cat. #11058021), and incubated for 20 min. The lipid complex was added to each well and incubated for 16 h. Following incubation, the medium was replaced with full serum medium. Cells were harvested and analyzed via flow cytometry at 36 h post-transfection following the method described above for quantifying EGFP knockout. Flow cytometry data was collected using LSRFortessa X-20 (BD Biosciences). EGFP and mCherry were excited using 488 nm and 561 nm lasers, respectively. Forward and side scatter was applied for doublet discrimination. Data was analyzed using FCSExpress (Dotmatics) by gating based on the EGFP fluorescence intensity relative to the negative control.

### Cryo-EM of Cas9 ternary complexes with chemically modified crRNA

Cas9–RNA–DNA ternary complexes were assembled at room temperature for 15 min in 20 mM Tris (pH 7.5), 200 mM KCl, 0.5 mM DTT, and 5 mM EDTA. Complexes (1.4 mg/mL) were applied to glow-discharged Quantifoil 2/2 holey grids and vitrified using an FEI Vitrobot Mark IV. Grids were inspected on a 200-kEv Glacios equipped with a Falcon IV direct electron detector and preliminary data sets were collected for 2 hours for each grid with acceptable density of particles and lacking visible aggregates. Final data sets were collected on a 300-kEv Titan Krios G3 cryo-TEM equipped with a Gatan K3 detector. Images were processed in cryoSPARC v4.1.2, including motion correction, CTF estimation, and particle selection using template-based picking and Topaz training, followed by 2D/3D classification and non-uniform refinement. Initial data included 4602 movies and yielded closed conformation structure at 3.7 Å. Additional 3492 movies were collected to improve resolution and to acquire a better representation of alternative 2D classes corresponding to the open conformation. Final resolution of the active closed conformation was 3.44 Å calculated with 73855 particles. The model was refined using Refmac program implemented in Phenix program suit with manual modeling using Coot. The refined structure and data were deposited with PDB ID: 9OTT and EMDB entry ID EMD-70859.

### Molecular dynamics simulations of Cas9 ternary complexes

The Cas9 dual guide complex (PDB ID: 8FZT) with both target strand (TS) and non-target strand (NTS) was used as the template for the Gaussian-accelerated molecular dynamics (GaMD) simulations. The missing residues in the protein template were filled using the loop modelling module in the UCSF chimera. The sequences of the nucleic acid components were altered to match experimental sequences using the “swapna” command in the UCSF chimera. The DNA modification was included at the 14^th^ position (which is the canonical 15^th^ position since we used 19 nt spacers) of the crRNA using X3DNA. The modeled protein and nucleic acid components were combined in the tleap module of AmberTools 22 to generate the modified (Mod) and unmodified (Unmod) complexes. The coordinates of the catalytic Mg^2+^ ions were included using the reported structure with PDB ID: 5Y36.7 The complexes were solvated using TIP3P water box with a dimension of 10 Å. A salt concentration of 100 mM NaCl was maintained.

The reported procedure for Cas9 complexes was used with slight modifications ^59^. Briefly, both complexes were subjected to energy minimization keeping the protein, RNA, DNA, and Mg^2+^ ions fixed with harmonic position restraints of 300 kcal/mol Å^2^. This has been carried out using the steepest descent method with 10,000 steps. Further, in the second stage, all the restraints were removed, and a similar steepest descent minimization was performed. Then, the complexes were heated from 0 to 50 K and from 50 to 100 K by running two NVT simulations of 50 ps each, imposing restraints of 100 kcal/mol Å^2^ on the protein, RNA, DNA, and Mg^2+^ ions. The temperature was then increased to 200 K in 100 ps of MD using an NVT ensemble, which reduced the restraint to 25 kcal/mol Å^2^. Finally, the system temperature was raised to 300 K in an NPT simulation of 500 ps without restraints. 1 ns NPT equilibration was done on the complexes, followed by a test production run of 10 ns NPT. After observing the trajectory of the system, production runs were carried out for 100 ns in the GPU-accelerated version of PMEMD ^60, 61^ in AMBER ^62^. SHAKE algorithm was applied to restrain bonds containing hydrogen with a default tolerance value of 0.00001 Å. The integration time step of simulations is 2 fs. Hydrogen atoms were added and were constrained to their equilibrium position with the SHAKE algorithm, and the temperature was maintained using Langevin ^63^ dynamics with a collision frequency γ = 1. Coupling the system to a Berendsen ^64^ barostat at a reference pressure of 1 atm and with a relaxation time of 2 ps gave pressure control. All the stages up to equilibration were performed using the SANDER module.

After 100 ns of conventional MD, GaMD simulations were performed for 500 ns with the potential being updated every 1.6 ns. 88 ns of NVT GAMD equilibration was done before the GAMD production. A dual boost protocol with the potential being applied to both the total energy and dihedral energy has been applied. The sigma0P and sigma0D parameters were kept at 6.0 and the k0P/k0D values were 1 at the end of the GaMD equilibration. The integration timestep, SHAKE tolerance, and the barostat and thermostat parameters were kept identical to the classical MD protocol.

Initially, the trajectory weights were extracted from the GaMD log files, where the energy values were given. Following this, the PMF is calculated upon accurate reweighting of the simulations using cumulant expansion to the 2^nd^ order. Reweighting was carried out using the PyReweighting toolkit by Miao and colleagues ^65^. Parameters like “Cut-off” and “disc” were altered to match our calculations. The PMF calculations were performed on the GAMD trajectories using amino acid distances and interdomain angles, which were previously determined experimentally, and can characterize the catalytic state of the protein. The convergence of the simulation was checked by calculating the PMF plots using varying bin sizes. Emax and cut-off values for reweighting were 20 and 10, respectively, for most calculations.

All the general analyses were performed using the CPPTRAJ ^66^ module of AmberTools 19 and VMD ^67^. For all the trajectories, the RMSD and the RMSF analyses are performed using the first frame of the simulation as the reference. The backbone RMSD of protein is calculated using the C, Cα, and N atoms, while the RMSD of the DNA/RNA includes the heavy atoms of the base and the sugar. RMSF calculations are calculated considering all the frames. The keywords “rms” and “atomicfluct” in the CPPTRAJ module have been used for these analyses. Distance analysis has been performed using the “distance” keyword. The distance between the specific atoms or the centroid of the whole residues was calculated. The “angle” keywords have been used for the respective angle calculations. The whole residues, range of residues, or specific atoms have been utilized for various calculations. Native contact is analyzed using the “nativecontacts” keyword. The native contacts were defined by the convention that every residue present within 5 Å of the reference residue during the first simulation frame is considered native contacts. The contacts formed during the simulation were considered non-native contacts. H-bond analysis was carried out in both CPPTRAJ and VMD. A cut-off of 135 degrees and 3.5 Å have been used in CPPTRAJ.

For cluster analysis, all the trajectories were clustered into 5 ensembles. The hierarchical agglomerative algorithm was utilized for the clustering. Every 10^th^ frame was considered for the clustering, and a sieve value of 10 was used for the clustering. The deviation in the whole protein was used as the metric for clustering. The major cluster representative structures were visualized in the PyMOL visualization software. This analysis was also performed in the CPPTRAJ module. The structural alignments were all carried out using the combinatorial extension algorithm in PyMOL software.

### Serum stability assays for quantifying crRNA nuclease resistance

3 µL of crRNA (300 pmol) in RNA resuspension buffer (10 mM Tris, pH 7, 0.1 mM EDTA) was combined with 5 µL of 10x PBS and brought to 25 µL with double-distilled water (ddH_2_O) at RT. To initiate the reaction, one vol (25 µL) of 100% human serum (Sigma Aldrich, Cat. #H4522), was added, mixed rapidly by pipetting, and incubated at 37°C for the indicated time. Reactions were stopped by addition of 25 µL of acid phenol (pH 4.5), vortexing for 15 s, then placing on ice. One vol (50 µL) of chloroform was added, samples vortexed for 15 s, then separated by centrifugation at 10,000 x g for 10 min at RT. The top aqueous layer was moved to a new tube, one vol (50 µL) of ddH_2_O added, sample vortexed for 30s, then placed at -20°C for 30 min. Samples were separated again by centrifugation and the aqueous layer removed and combined with the first aqueous layer that was extracted. Sample was precipitated by addition of 9 vols (900 µL) of 2% LiClO_4_ in acetone, vortexing for 30 s, and incubation at -20C. Precipitated product was separated by centrifugation at 10,000 x g for 15 min, the pellet washed gently with acetone and centrifuged again for 5 min, then air dried at RT.

Dried RNA pellets were resuspended in 1x denaturing loading buffer (90% formamide, 1x TBE), denatured at 95°C for 3 min, then resolved on a denaturing 15% urea polyacrylamide gel (7M urea, 1x TBE, 15% 19:1 acrylamide:bisacrylamide, 1% glycerol). Bands were visualized by methylene blue staining and destaining in water and gels imaged using a ChemiDoc MP Imaging System (BioRad). Degradation of crRNA was quantified using ImageJ. The intensity of the intact, full length crRNA band divided by the combined intensity of all bands in the lane for each sample was taken as the fraction of intact full-length crRNA that remained.

### On Target and Off Target Editing Quantified by Nanopore Amplicon Sequencing

HEK293T cells stably expressing EGFP and SpCas9 were kindly provided by Dr. Wen Xue (UMass Chan Medical School) ^57^. To increase SpCas9 expression, 2 x 10^6^ cells were electroporated with 9 μg of LentiV_Cas9_Puro (Addgene #108100) using Neon NxT electroporation system (ThermoFisher Scientific) and the 100 μL Neon Transfection System Kit (ThermoFisher Scientific, Cat. #MPK10096) using a program of 1,100 V, 20 ms per pulse, 2 pulses. After 48 h, 85,000 cells were reverse-transfected in a 48-well plate, with 40 pmol of 1:1 molar ratio crRNA:tracrRNA duplex using 1 µL Lipofectamine RNAiMAX (ThermoFisher Scientific, Cat. #MPK10096), diluted in Opti-MEM (ThermoFisher Scientific, Cat. #11058021) and incubated for 12 h. Following incubation, the medium was replaced with full serum medium and harvested after 24 h for genomic DNA extraction using Monarch Genomic DNA purification kit (NEB, Cat. #T3010L) following the manufacturer’s recommended protocol. DNA was quantified by absorbance at 260 nm.

For each sample, multiplex PCR was carried out using pooled primer sets that generate ∼1 kb amplicons centered on either the intended Cas9 cut site (on-target) or one of the validated off-target sites previously reported ^68, 69^. Reactions (25 µL) contained 30 ng of genomic DNA and were amplified with Q5 Hot Start High-Fidelity DNA Polymerase (NEB, M0493L) following the manufacturer’s protocol. The PCR program was set to 98°C for 3 min followed by 30 cycles of 98°C for 10 sec, 66°C for 30 sec, 72°C for 45 sec, and a final extension of 72°C for 2 min. Primer sequences and final concentrations of each primer can be found in **Table S2**.

Library preparation of amplicons was performed using Native Barcoding Kit 96 V14 (SQK-NBD114.96) (Oxford Nanopore Technologies, ONT) according to the manufacturer’s protocol. Briefly, PCR amplicons were concentrated and purified via SPRI bead cleanup. Amplicons were end-prepped using 0.75 μL Ultra II End-prep Enzyme mix, and 1.75 μL Ultra II End-prep reaction buffer (NEB) and incubated at 20°C for 10 min followed by 65°C for 10 min using a thermal cycler. Reaction clean-up was performed using a 1:1 volume ratio of reaction to AMPure XP beads (Beckman Coulter) following the manufacturer’s recommended protocol and then eluted. The amplicons were then individually barcoded. Briefly, 7.5 μL of the end-prepped DNA, 2.5 μL of barcode, 10 μL of Blunt/TA Ligase Master Mix (NEB) were combined and incubated at room temperature for 20 min. The reactions were stopped by adding 2 μL of 500 mM EDTA to each barcoding reaction and pooled together. The pooled samples were purified using AMPure XP beads (Beckman Coulter) and eluted in 30 μL of elution buffer (10 mM Tris-HCl, pH 8.5). The eluted barcoded samples were adapter ligated by combining 30 μL of pooled barcoded samples, 5 μL of native adapter (NA), 10 μL of 5x NEBNext Quick Ligation Buffer (NEB), 5 μL of Quick T4 DNA Ligase (NEB) and incubated for 30 min at RT. The library was purified using AMPure XP beads, loaded on a PromethION R10.4.1 DNA flow cell and sequenced on an Oxford Nanopore Technologies (ONT) P2 solo using MinKNOW. Live coverage visualization was established using RAMPART (https://github.com/artic-network/rampart).

On-target and off-target editing frequencies were quantified with CRISPResso2 (https://github.com/pinellolab/CRISPResso2) using whole-genome-sequencing (WGS) parameters for Cas9. For each sample, background signal was removed by subtracting the editing level observed in samples where no crRNA was included during treatments.

## Results

### Ribose replacements at crRNA-Cas9 2′-OH contact residues inhibit gene editing

Previous investigations of Cas9 guide RNA modification could not achieve complete replacement of all nucleotides with ribose sugar modifications, which was later linked to several specific residue positions ^33, 38, 42–45^. Structural inspection indicated that the 2’-OH group of these guide RNA residues appear to make polar contacts, and therefore hydrogen bond, with the Cas9 peptide backbone ^46, 70^, suggesting a critical but unknown role for these contacts in catalysis. To assess the role of the 2’-OH at these positions, we synthesized a series of crRNAs with a 19 nt spacer sequence targeting an EGFP reporter gene (**Table S1**). These crRNAs incorporated 2’-arabinonucleic acid (2’-ara) or 2’-fluoro (2’-F) substitutions at two or more of the six crRNA 2’-OH contact positions while all other residues were held constant as either RNA or 2’-F. (**Fig. 1A**). The 2’-ara is a stereoisomer of RNA, where the 2’-OH is maintained but positioned on the top (beta) face of the ribose, inducing a DNA-like C2’/O4’-*endo* (South/East) sugar pucker ^56, 71, 72^. In contrast, 2’-F adopts a strong RNA-like C3’-*endo* (North) sugar pucker ^73^ while sacrificing hydrogen bond donor capacity. We reasoned that these two modifications would allow us to probe hydrogen bonding and sugar conformation. We evaluated activity via *in vitro* cleavage assays and cell-based EGFP knockout (KO) assays (**Fig. 1A**). For several modification schemes, such partial replacements with 2’-ara (crEG-SJ2ara and crEG-SJ7) or an all-2-’F design (crEG-SJ16), we observed high *in vitro* cleavage activity but little or no cell-based EGFP KO activity. In contrast, only one modification scheme, crEG-SJ3, which was comprised of all 2’-F but retained RNA at the six critical residue positions, provided full rescue of cell-based editing. Substituting the last two critical contact positions of crEG-SJ3 with 2′-ara resulted in complete loss of cell-based editing (crEG-SJ28), underscoring the sensitivity of individual positions. Together, these results confirmed the dependence of Cas9 gene editing on the six residues predicted to make 2’ polar contacts with the Cas9 backbone. The discrepancy between *in vitro* and cell-based assays also indicated that gene editing is particularly sensitive to sugar modification, perhaps due to the cellular environment.

**Fig. 1.**
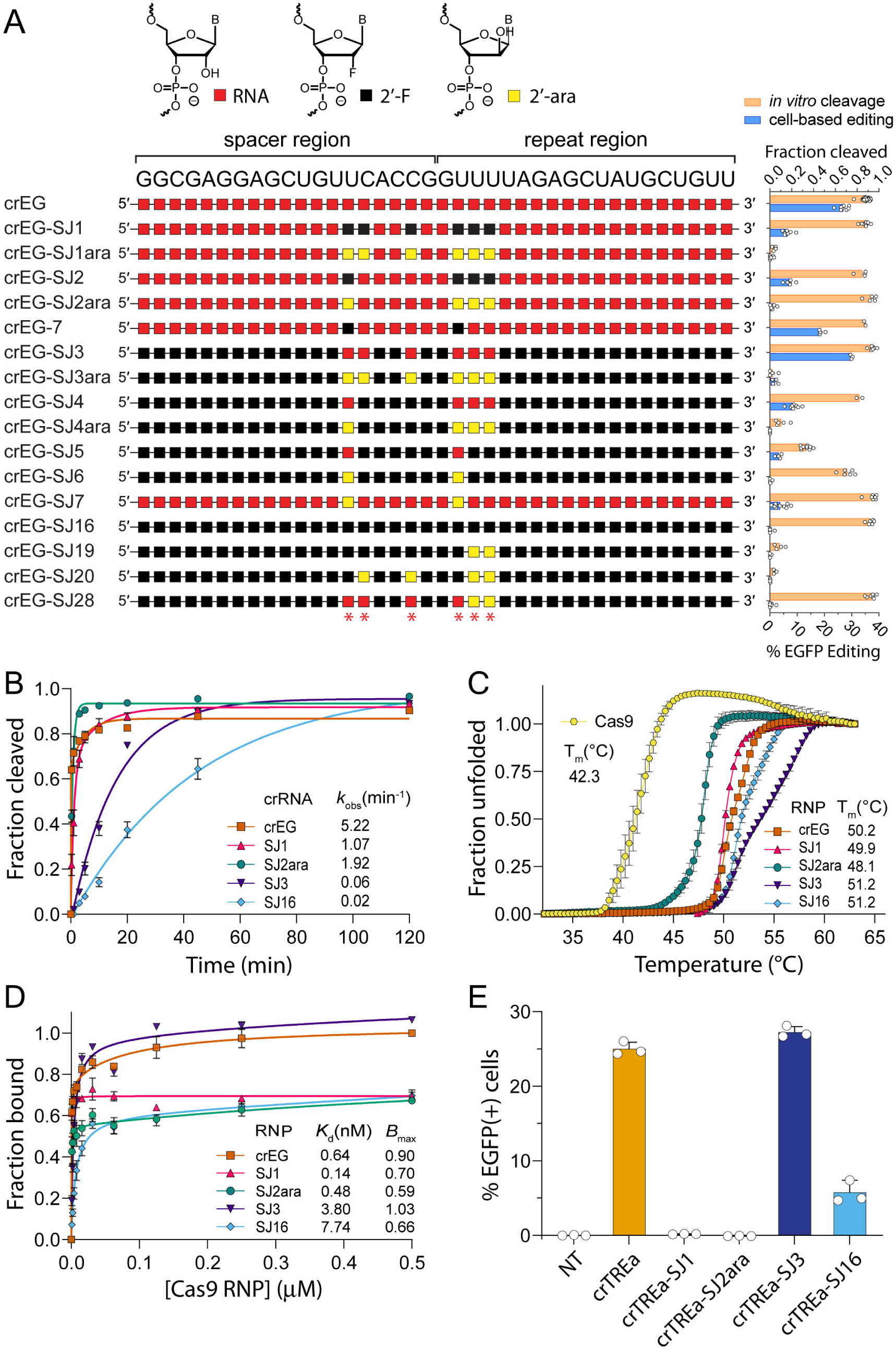
CRISPR-Cas9 editing and target engagement are mediated by crRNA 2’-hydroxyl contacts. (**A**) Schematic of the crRNA sequence used for targeting an EGFP reporter gene and chemical modifications used to probe 2’-OH contacts. *In vitro* cleavage activity (orange) and cell-based EGFP editing (blue) is shown on the right. 2’-OH contacts with SpCas9 are indicated with red asterisks below. n = 3 or more experimental replicates. Error bars are S.E.M. (**B**) Time-course of *in vitro* cleavage activity using select crRNAs from panel A. Curves were fitted to an exponential two-phase decay equation. Error is reported as S.E.M. (**C**) Thermal denaturation of Cas9 RNP complexes assayed by absorbance at 280 nm to measure melting temperature (*T*_m_). n = 2 experimental replicates. Error bars are S.E.M. (**D**) Target DNA binding by dCas9 RNP measured by dot blot filter binding of radiolabeled target DNA. Curves were fit to a one-site binding curve. n = 2 experimental replicates. Error bars are S.E.M. (**E**) CRISPRa-based assay to measure dCas9-VPR binding guided by crTREa crRNA to a Tet-On 3G promoter driving EGFP in HeLa cells. Unmodified crTREa is shown as a control. EGFP expression was quantified by flow cytometry. N = 3 experimental replicates, Error is S.E.M.

### Editing efficiency correlates with DNA target engagement

We hypothesized that ribose replacement at critical residues may reduce catalytic rates, compromise RNP assembly, or restrict target DNA engagement. To investigate these possible biochemical explanations, we systematically tested several representative modified crRNAs for their effects on *in vitro* cleavage kinetics, ribonucleoprotein (RNP) stability, and DNA binding. Selected crRNAs were crEG, crEG-SJ1, crEG-SJ2ara, crEG-SJ3 and crEG-SJ16. These were chosen for their opposing or complementary chemical substitutions and activities during *in vitro* cleavage and cell-based editing (**Fig. 1A**).

We first conducted *in vitro* time course cleavage assays, hypothesizing that slow cleavage kinetics may contribute to reduced cell-based editing (**Fig. 1B**). Interestingly, crEG-SJ2ara, which did not support cell-based editing, exhibited the second highest *k*_obs_ value of 1.92 min^-1^. In contrast, crEG-SJ3, which conferred the highest cell-based editing activity, showed slower kinetics, with a *k*_obs_ of 0.06 min^-1^ (**Fig. 1B**). These results did not support cleavage kinetics as an explanation for cell-based editing efficiency, as evidenced by the lack of a significant correlation (*r* = 0.32) (**Fig. S1A**).

Inefficient formation of Cas9-crRNA-tracrRNA complexes, an essential first step, could compromise catalysis. Reduced RNP stability might help explain a loss in cell-based editing since cells may have limited concentrations of RNP compared to *in vitro* cleavage conditions. We therefore performed UV-monitored (absorbance at 280 nm) thermal denaturation analysis to quantify global RNP stability (**Fig. 1C**). We found that crEG-SJ3, which exhibits high cell-based editing, had a melting temperature (*T*_m_) value of 51.2 ± 0.3°C. However, crEG-SJ16, which showed complete loss of cell-based editing, had the same *T*_m_ value. These two contrasting examples illustrate that RNP stability is unlikely to be the primary determinant of reduced cell-based editing, as reflected by the lack of a significant correlation (*r* = 0.41) (**Fig. S1B**).

To evaluate target DNA binding, we used a dot blot filter binding assay and catalytically “dead” Cas9 (dCas9) containing D10A and H840A mutations ^74^. Increasing concentrations of dCas9, in complex with a modified candidate crRNA and unmodified tracrRNA, was titrated onto a radiolabeled double-stranded DNA target substrate and binding quantified by dot-blot analyses (**Fig. 1D**). Relative binding was determined by fitting to a one-site binding hyperbola and calculating the apparent dissociation constant (*K*_d_) and maximal binding capacity (*B*_max_), which reflects the fraction of available DNA sites bound at saturation. Unmodified crEG exhibited a high binding affinity (*K*_d_ = 0.64 nM) and binding capacity (*B*_max_ = 0.90) while modified crRNAs that supported little or no cell-based editing, specifically crEG-SJ1, crEG-SJ2ara, and crEG-SJ16, all showed relatively strong binding affinities but reduced *B*_max_ values of 0.70, 0.59, and 0.66, respectively. Conversely, crEG-SJ3, possessed a high *B*_max_ of 1.03. We observed no significant correlation between cell-based editing activity and *K*_d_ (*r* = -0.18) (**Fig. S1C**), while a strong positive correlation was found for *B*_max_ (*r* = 0.95, *p* = 0.01) (**Fig. S1D).** These results suggest that ribose replacement at 2’-OH contact residues in crRNAs can increase the fraction of Cas9 RNP complexes incompetent for target binding while maintaining high binding affinity for competent complexes that remain.

To determine whether reduced target DNA binding was also observed inside cells, we developed a CRISPR activation (CRISPRa) assay. In this system, an EGFP expression cassette driven by a tetracycline-inducible (Tet-On) promoter is stably integrated into HeLa Tet-On 3G cells. A vector expressing dCas9-VPR, the CRISPRa protein, is electroporated into these cells, followed by transfection of tracrRNA and a crRNA (crTREa) that targets seven tandem tetracycline receptor sequence elements in the Tet-On promoter. Binding of multiple dCas9-VPR proteins, guided by crTREa, will induce transcription of EGFP. Higher fluorescence indicates stronger dCas9-VPR binding, allowing us to infer relative binding efficiency from EGFP expression levels. With this model system, we tested the same modification patterns used above in the crTREa guide and measured EGFP by flow cytometry (**Fig. 1E**). EGFP activation trends mirrored *B*_max_ results from filter binding assays, with only crTREa-SJ3 showing strong activation of EGFP, similar to the unmodified crTREa control. A positive correlation was observed between CRISPRa results and cell-based editing efficiency (*r* = 0.95, *p* = 0.015) (**Fig. S1E**). These results indicate that reduced target engagement in cells best correlates with loss of gene editing due to ribose sugar modification at critical 2’-OH contact positions ^75^.

### 2’-F ribose replacement induces alternative Cas9 ternary complex conformations

To investigate potential structural effects of ribose replacement at crRNA 2’-OH contacts, we performed cryogenic electron microscopy (cryo-EM) on a Cas9 ternary complex assembled with unmodified tracrRNA and crEG-SJ1, which contains 2’-F at the six contact residues (**Fig. 2A**), as well as a complementary double-stranded DNA target similar to one previously reported for structural studies ^46^. EDTA was included to prevent divalent metal ion-dependent cleavage of the target. Initial 2D classification of crEG-SJ1 ternary complexes resulted in four distinct classes. We observed a major “closed” conformation for ∼50% of all particles while ∼1/3 fell into two “open” conformations, one with density for half of the structure and one resembling a previously published open state ^76^. The remaining small percentage of particles were classified into a poorly structured “distorted” class (**Fig. 2B**) that was not observed in cryo-EM data for crE2, an unmodified dual-guide RNA (dgRNA) ternary complex (**Fig. 2C**) ^77^. These 2D class distributions support the hypothesis that modified crRNAs may induce formation of relatively stable alternative complexes that are not competent for target binding or cleavage.

**Fig. 2.**
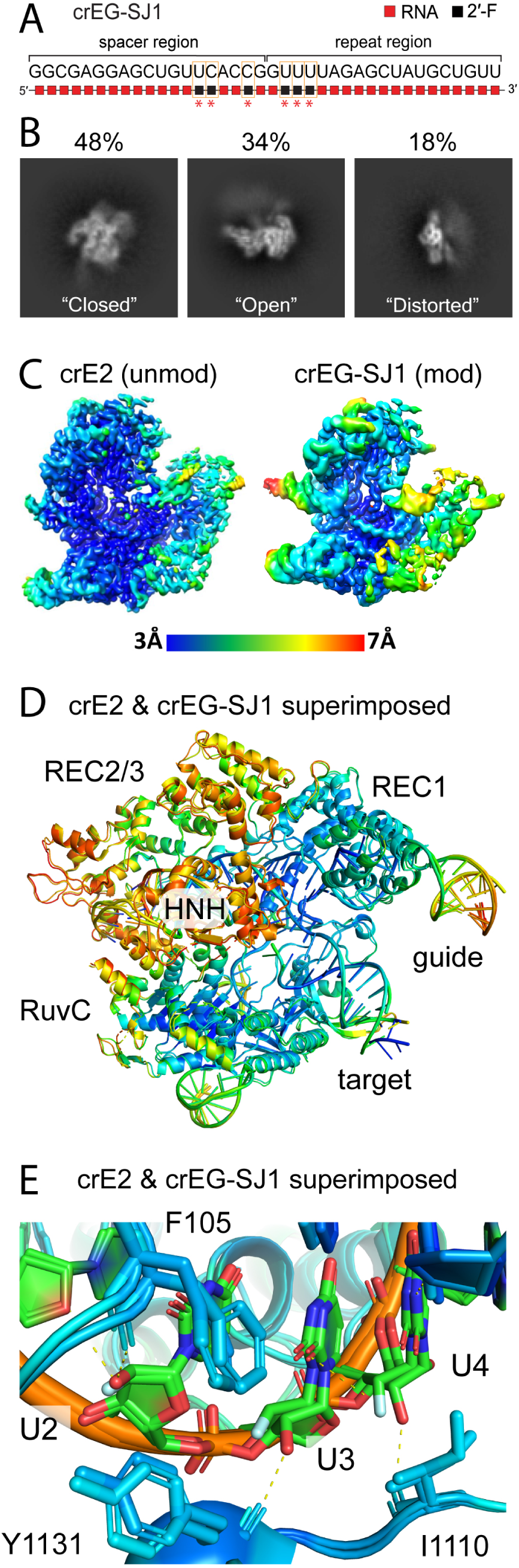
Cryo-EM analysis of a Cas9 ternary complexes assembled with crEG-SJ1 indicates conformational heterogeneity. (**A**) Sequence and modification scheme for crEG-SJ1 crRNA used for structural investigations. (**B**) Representative images of closed, open, and distorted 2D classifications for Cas9-crEG-SJ1-tracrRNA-DNA (ternary) complexes. (**C**) Electron density and resolution of the 3D reconstruction for Cas9 assembled with unmodified crE2 crRNA and modified (crEG-SJ1) crRNA. The coloring scheme represents the resolution within each structure, ranging from 3 Å to 7 Å. (**D**) Superimposed reconstructed 3D structure of Cas9 ternary complex assembled with an unmodified (crE2) and modified (crEG-SJ1) color coded by B-factor, with blue being highest and red lowest, following 3D reconstruction. (**E**) Close-up of uridine residues modified to 2′-F in crEG-SJ1 in the superimposition shown in panel E. The amino acids that make backbone contacts with the critical 2′-OH contacts are indicated, specifically F105, I1110, and Y1131. Hydrogen bonds are indicated by dotted yellow lines. Fluorine is shown as blue and hydroxyl is shown as red on residues U2-U3.

Although we could not refine the open or unstructured classes into 3D reconstructions with resolutions below 7.2 Å, the closed conformation yielded a final 3D reconstruction at 3.44 Å (**Fig. 2C**). We compared this structure to a high-resolution reconstruction of the unmodified crE2 ternary complex (PDB ID: 8FZT) that we previously resolved in a separate study ^77^. Global structures for crE2 dgRNA and crEG-SJ1 were overall remarkably similar in conformation (**Fig. 2D**, **S2A-B**), although increased flexibility was found in the REC2/3 and HNH domains (**Fig. S2C-D**). The 2’-F substituted residues assumed conformations in the closed structure nearly identical to the RNA residues in the unmodified crE2 ternary structure (**Fig. 2E**). Likewise, the Cas9 peptide backbone and amino acid side chains of F105, I1110, and Y1131, where 2’-OH contacts are predicted in native structures, were essentially unchanged (**Fig. 2E**). These results indicate that crEG-SJ1 supports closed conformations similar to a native, unmodified ternary complex but also induces the stabilization of alternative structures.

### Specific 2’-OH contact residues restrict gene editing and target engagement in cells

To determine the contribution of individual 2’-OH contact residues on Cas9 editing, we walked several ribose replacement chemistries individually across each position and quantified their impact on EGFP KO in HEK 293T cells (**Fig. 3A**). Chemical modifications were designed to either retain some degree of H-bonding capacity, such as 2’-F ^78^ and locked nucleic acid (LNA) ^79^ (H-bond acceptors), 2’-amino (2’-NH_2_) ^80^, 2’-amino locked nucleic acid (2’-NH-LNA) ^81^, and unlocked nucleic acid (UNA) ^82^, or to eliminate H-bonding potential, such as 2’-H (DNA), and 2’-*O*-methyl (2’-OMe) ^83^. Modifications like 2’-F, 2’-NH-LNA, LNA, and 2’-OMe also induce strong RNA-like sugar pucker while DNA and UNA introduce significant flexibility ^84^. Despite the chemistry employed, a general trend was observed across all six probed positions (**Fig. 3B**). The central residue 4, and to a lesser extent 3 and 5, were the most tolerant of modification while terminal residues, 1, 2, and 6, were more sensitive. These results suggest an intrinsic role for these residues in regulating Cas9 editing. Some ribose replacement chemistries performed better at certain positions. For example, position 4 tolerated DNA, LNA, and 2’-NH-LNA very well; positions 3 and 5 tolerated DNA and 2’-F the most; 2’-F performed best at position 1; and position 6 best tolerated 2’-F and 2’-NH-LNA. Modification resulted in reduced gene editing for all residues except position 4. To assess sequence-specific effects, we designed a second crRNA, crEG_2, targeting a different sequence of the EGFP reporter. We repeated the single modification walk using DNA and found that editing followed a similar trend as before, with positions 3 and 4 tolerating modification the most, while positions 1, 2, and 6 were the least tolerant (**Fig. 3C**).

**Fig. 3.**
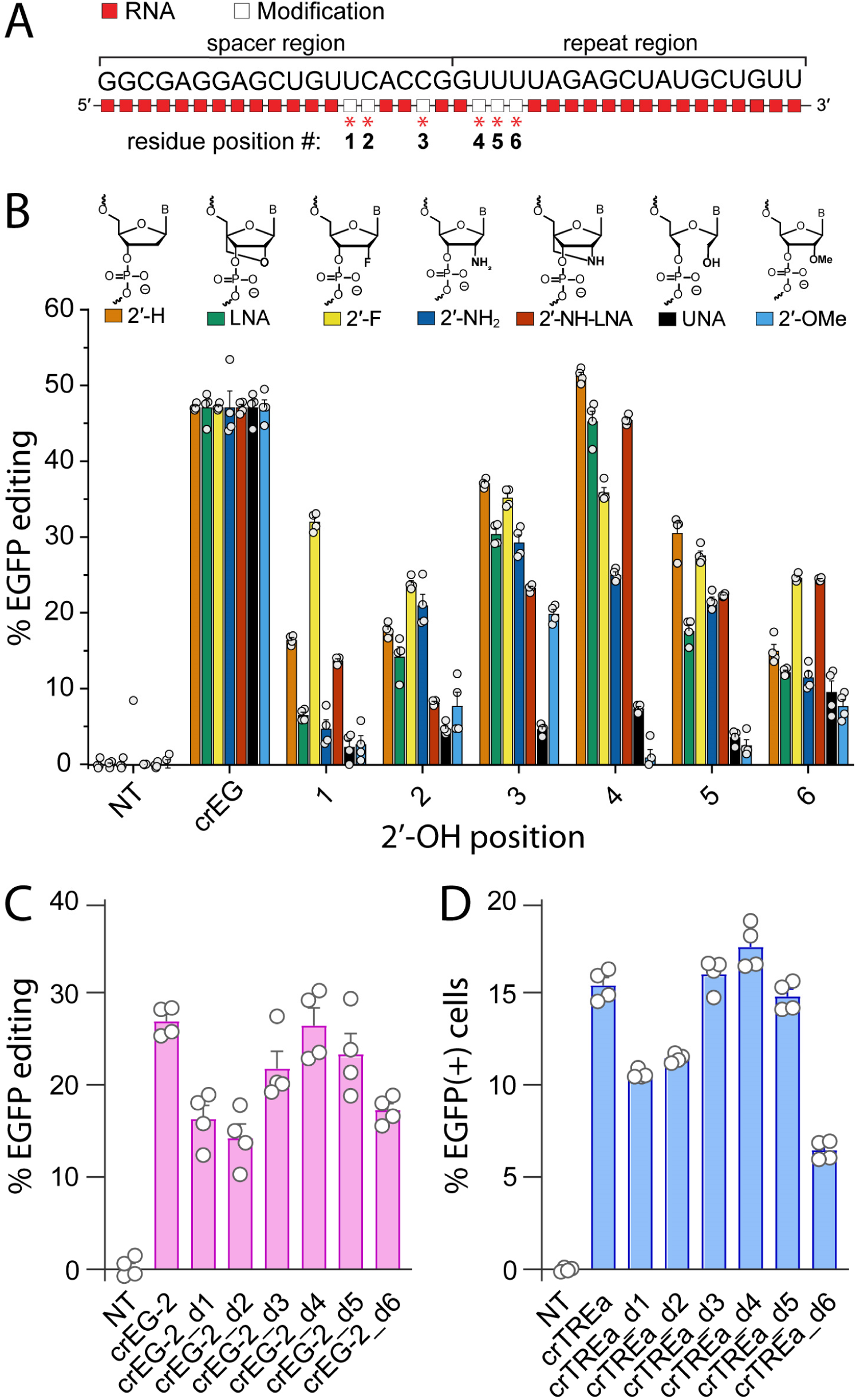
Individual 2’-hydroxyl contacts differentially tolerate chemical modification and influence Cas9 gene editing and DNA binding in cells. (**A**) Illustration of the six 2’-OH contact positions screened individually by chemical modification. (**B**) Chemical structures of ribose modifications tested. Cell-based editing of EGFP using crRNAs with modifications individually “walked” across each 2’-OH contact position. n = 4 experimental replicates. Error bars are S.E.M. (**C**) Cell-based editing of EGFP using crEG-2 targeting an alternative sequence in EGFP. A single DNA modification was walked across each 2′-OH contact position. n = 4 experimental replicates. Error bars are S.E.M. (**D**) CRISPRa of crRNAs with a single DNA modification walked across each 2′-OH contact position. EGFP expression was quantified by flow cytometry. n = 4 experimental replicates. Error bars are S.E.M.

To determine if position-specific ribose replacement was correlated with target binding in cells, we performed CRISPRa using crRNAs where DNA was walked through each 2’-OH contact residue (**Fig. 3D**). The trend in EGFP activation was similar to the trend of cell-based editing. Consistent with gene editing results, activation of EGFP expression was highest when DNA was substituted at position 4 (crTREa_d4). In contrast, DNA at positions 1, 2, and 6 resulted in the lowest activation of EGFP expression. The effect on DNA target binding at each position positively and significantly correlated with cell-based editing (*r* = 0.89, *p* = 0.008) (**Fig. S1F**). These findings support that 2’-OH contact residues are crucial for Cas9-mediated target DNA binding and catalysis, both individually and in combination.

### 2’-OH contacts with Cas9 regulate conformational dynamics essential for target engagement

We hypothesized that the distribution of 2D particle classes during cryo-EM may be the result of complexes partitioning into competent and non-competent states. To investigate the mechanism underlying potential conformational and allosteric effects, we performed 1 μs Gaussian-accelerated molecular dynamics (GaMD) simulations. We substituted a single DNA in place of RNA at the 2’-OH contact position 1 (spacer nt 14) and refer to this as the crEG_d1 Mod complex (**Fig 4A**). A matched unmodified complex with crEG (Unmod) was also generated. Protein, RNA, and DNA (target strand) backbone RMSD values revealed no sudden changes, indicating well-equilibrated simulations (**Fig. S3**). The Cα distances between residues S867 (HNH) and N1054 (RuvC), which have been associated previously with the apo (6 Å), RNA bound (7 Å), DNA bound (28 Å), pre-catalytic (45 Å), and catalytic states (57 Å) of Cas9 ^85^, were then calculated (**Fig. S4**). The distance plot revealed that the Unmod complex retains the DNA-bound state (∼28-31 Å) while the Mod complex assumes an inactive apo-like conformation (∼6-7 Å) (**Fig. 4B**). A reduction in the catalytic state was also supported by increased distance measured between the catalytic H840 residue of the HNH domain and the DNA scissile phosphate ^86^ in the Mod complex (**Fig. 4C**).

**Fig. 4.**
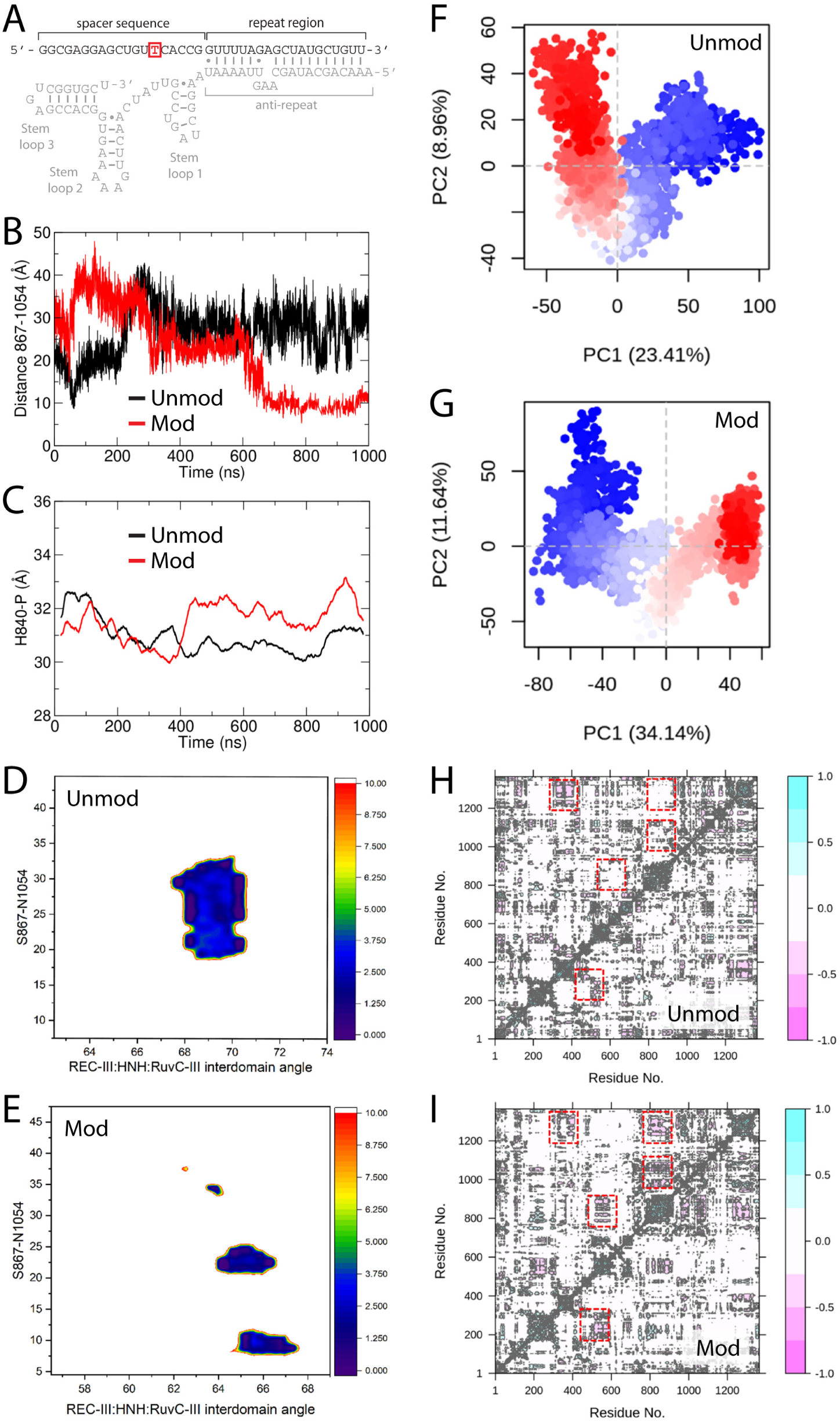
Molecular dynamics simulations reveal major conformational shifts in the Cas9 ternary complex toward an inactive state when a single 2’-OH contact is lost. (**A**) Secondary structure of the crEG guide RNA with the single DNA nucleotide substitution in crEG_d1 highlighted in red within the spacer. (**B**) Time evolution of the distance between residues S867 and N1054 in the REC3 domain over 1 μs simulations. Complexes with unmodified crEG (black) or DNA-modified crEG_d1 (red) show larger fluctuations in the modified complex. (**C**) Distance between the catalytic HNH residue H840 and the scissile phosphate of the target strand for unmodified crEG (black) and modified crEG_d1 (red). (**D**, **E**) Two-dimensional potential of mean force (PMF) landscapes showing the distribution of interdomain angles between REC3, HNH, and RuvC for complexes with unmodified crEG (**D**) or modified crEG_d1 (**E**). (**F**, **G**) Principal component analysis of Cas9 motions. Each point represents an instantaneous trajectory frame projected onto PC1 and PC2 for complexes with unmodified crEG (**F**) or modified crEG_d1 (**G**). Frames are colored from blue (early) to red (late) to illustrate conformational sampling shift over time. (**H, I**) Dynamic cross-correlation matrices of Cα atom motions for complexes with unmodified crEG (**H**) or modified crEG_d1 (**I**). Boxed regions highlight the loss of correlated motions in the crEG_d1 complex.

Widening of the groove accommodating the RNA:DNA hybrid is necessary target strand (TS) cleavage and conformational transition of the HNH domain.^3^ Therefore, interdomain angles between REC-III:HNH:RuvC-III and REC-II:REC-I:PI were sampled (**Fig. S5A**). While the angle between REC-II:REC-I:PI remained quite similar across complexes, the angle between REC-III:HNH:RuvC-III was smaller for the Mod complex (59-61°) versus that of the Unmod complex (69-71°) (**Fig. S5B-C)**. Thus, conformational changes in Cas9 at domains involved in accommodating the guide RNA-TS hybrid (R-loop) structure appear to be inhibited and insufficient in the Mod complex.

To better understand energy minima distribution, we constructed potential mean of force (PMF) plots using the C_α_ distance between S867-N1054 and the interdomain angles between REC-III:HNH:RuvC-III. The trajectories were subjected to a hierarchical clustering algorithm to generate 5 ensembles. While the Unmod complex generated only a single large population, the ensemble is rather scattered in the Mod complex (**Fig. S5D**). In addition, the Unmod complex showed primarily a single low-energy well (**Fig. 4D**) whereas multiple scattered low-energy wells were observed in the Mod complex (**Fig. 4E**). These results support formation of multiple conformations for the Mod complex, some of which occupy a catalytically incompetent state.

Superimposition of the major clusters of the Unmod and Mod complexes (**Fig. S6**) showed major changes in the overall complex. In all domains, large changes in the loop orientations were observed, helices and loops in the REC domains were modestly displaced, connecting loops between HNH and RuvC were altered, and significant changes were found in the RuvC domain.

Principal component (PC) analyses were carried out to investigate global impacts on Cas9 dynamics. With respect to time, the PC values show an opposite trend in the Unmod and Mod complexes, revealing that their dynamics are the reverse of one another (**Fig. 4F-G**). The PC values also exhibit a broader range in the Unmod complex, indicating that more conformational space is spanned by the Unmod complex. The final value of PC is negative in Unmod and positive in Mod, indicating that the Mod complex does not achieve a competent state since negative PC1 values are associated with the active closed state of Cas9 ^87^. To gain insight into Cas9 domain motions, we generated dynamic cross-correlation matrix (DCCM) plots, which revealed four regions where neutral or correlated motions became anticorrelated in the Mod complex (**Fig. 4H-I**).

Global changes in the Mod complex suggest that dynamic relationships within the Cas9 complex are altered. This would be expected to repartition the network distribution of amino acids and change allosteric signaling pathways. To probe these changes, the amino acid network was grouped into multiple communities (C) (**Fig. 5A-B**). In the Unmod complex, C1 consists almost entirely of bridge helix (BH), PAM-interacting (PI) domain, and most of the RuvC residues. C2 harbors residues from the BH, RuvC and REC-I domains, C4 contains most of the REC-III domain residues, C3 contains REC-II residues, and C5 consists primarily of the HNH domain. C2 and C3 have a strong connection, C4 has a strong connection with C1, C2, and C3, and C5 is well connected to all other communities, which fits with expected mechanisms. In contrast, the Mod complex displays a large redistribution in the community pattern with C1, C2, and C4 communities shrinking considerably and C3 and C5 communities growing. Importantly, canonical Unmod connectivity patterns between communities are lost and redistributed. C1 in the Mod complex now contains most of the HNH (formerly in C5) and REC-II (formerly in C3) residues and BH and PI domain residues are now found in C2 (formerly in C1). The Mod community with the HNH domain (C1) fails to establish strong connections and RuvC residues are scattered across communities. Alterations in the BH-REC-RuvC domain distribution most likely block communication between recognition and nuclease lobes.

**Fig 5.**
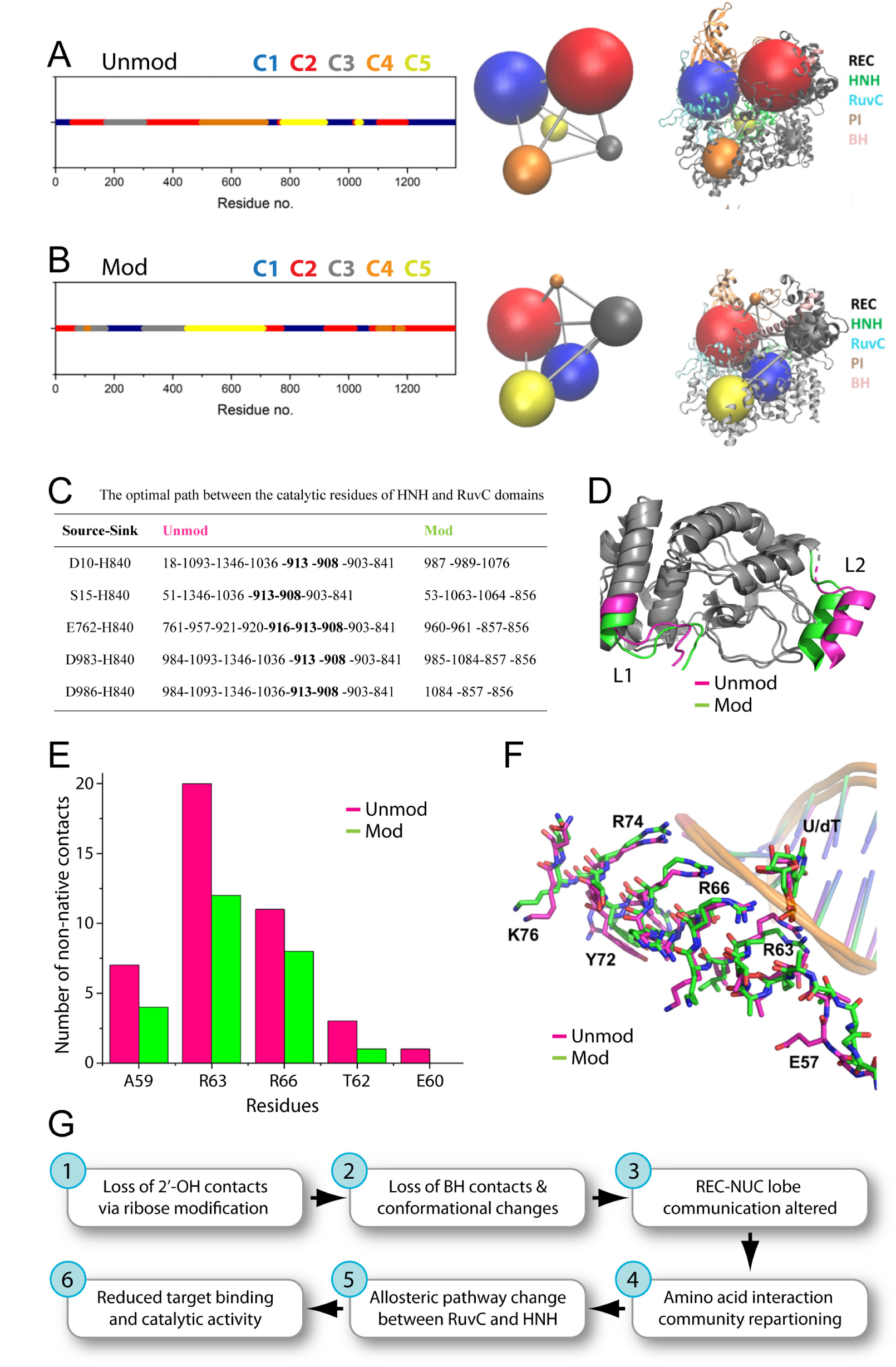
DNA substitution in the crEG_d1 crRNA disrupts Cas9 communication networks and target DNA contacts. (**A, B**) Dynamic network community analysis of Cas9 bound to (**A**) unmodified crEG or (**B**) modified crEG_d1. Linear maps show residue assignment to communities C1-C5 along the Cas9 sequence, and the corresponding three-dimensional representations show each community as a sphere with edges indicating intercommunity contacts mapped onto the Cas9 structure (REC, HNH, RuvC, PI, BH). (**C**) Optimal communication paths between the catalytic residues of the RuvC (D10) and HNH (H840) domains for unmodified crEG and modified crEG_d1 complexes. Residue sequences are listed from source to sink; bold entries highlight segments that differ between complexes. (**D**) Overlay of representative structures from the major clusters of unmodified (magenta) and modified (green) simulations showing altered orientations of loops L1 and L2 that flank the HNH active site. Catalytic domain helices are shown in gray. (**E**) Bar graph showing the number of non-native contacts formed between selected bridge helix residues (A59, R63, R66, T62, E60) and the target U/dT nucleotide in unmodified crEG (magenta) and modified crEG_d1 (green) complexes. (**F**) Superposition of major clusters illustrating the positions of bridge helix residues (K76, Y72, R74, R66, R63, E57) relative to the target U/dT base for unmodified (magenta) and modified (green) complexes. (**G**) Schematic model summarizing the mechanistic cascade that begins with loss of 2′-OH contacts (crEG_d1) and results in reduced target binding and catalytic activity.

To specifically assess changes in allosteric signaling pathways, with an emphasis on communication to HNH and RuvC ^88^, we calculated the optimal paths between catalytic residues D10, S15, E762, and H983 (**Fig. 5C**). Unlike in the Unmod complex, the paths of the Mod complex were much shorter and residues in the L2 loop, particularly 908, 913, and 916, were absent. The L2 loop, also known as linker 2, connects the HNH and RuvC domains and is an allosteric transducer ^88^ critical for the function of checkpoint mechanisms that regulate Cas9 target engagement and catalysis ^69, 89, 90^. Visualization of the orientation of these loops in major clusters shows significant deviations in the Mod complex (**Fig. 5D**).

The Cas9 domain closest to the site of DNA modification in the crRNA is the BH. Therefore, native and non-native contacts, which formed during the simulation, between the U/dT:dA base pair (crRNA:TS) were calculated. Native contacts to BH residues R63 and R66 were lost and fewer non-native contacts to A59, T62, R63, R66, and E60 were formed in the Mod complex (**Fig. 5E**). The orientation of BH residues was also altered in the Mod complex (**Fig. 5F**). Thus, incorporation of a single DNA at one 2’-OH contact position disrupts dynamic interactions with the BH, which is critical for the coordinated action of HNH and RuvC domains ^91^. Together, these results suggest that loss of only one 2’-OH interaction with the BH is sufficient to induce changes in the community network, inefficient communication between the REC, RuvC, and HNH domains, and rewiring of allosteric signaling required for efficient target DNA binding and catalysis (**Fig. 5G**).

### Modulation of Cas9 dynamics with phosphorothioate enables full ribose replacement

To take advantage of the mechanistic insight gained, we generated heavily and fully modified crRNAs using chemical modification schemes based on our results and previous literature ^44, 50, 70^. We began with a crRNA possessing eight 2’-OMe modifications and three phosphorothioate (PS) linkages at the termini and 2’-F at all internal positions except the six 2’-OH contact residues. To this control design (crEG-FMC), we added PS linkages 3’ of each 2’-OH contact residue or replaced the RNA with 2’-F or DNA based on previous modification walks. These designs, crEG-FM1 through crEG-FM4, whittled down the native RNA content while maximizing nuclease resistance and Cas9 compatibility. Indeed, these designs all performed as well or better than the unmodified crEG (**Fig. 6A**). To explore the importance of RNA and PS linkages at positions 2 and 6, we replaced one remaining RNA residue at a time with 2’-F and removed or added PS linkages (**Fig. 6A**). Each RNA residue played a critical role that individually reduced editing (crEG-FM4.2 and crEG-FM4.3). PS linkages, in contrast, did not alter editing efficiency in this context (crEG-FM4.1 and crEG-FM4.4). These results underscore the critical nature of the individual RNA residues at positions 2 and 6. Despite possessing 3’ PS protection at the two remaining RNA residues, incubation crEG-FM4 in 50% human serum resulted in rapid degradation (**Fig. S7**).

**Fig 6.**
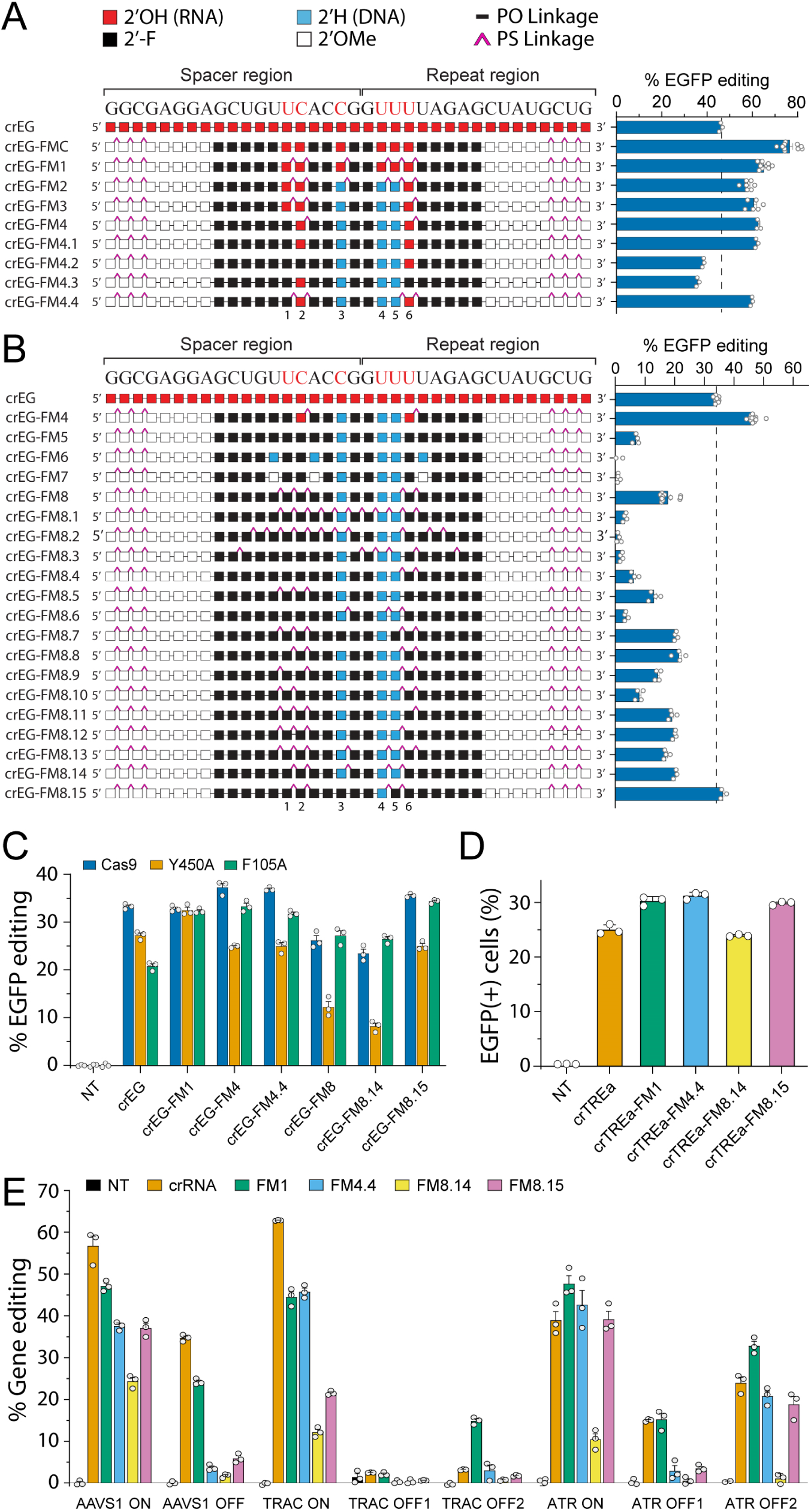
Ribose free crRNA preserves target engagement, supports on-target editing, and reduces off-target activity. **(A)** Cell-based EGFP editing using heavily modified crRNAs toward full ribose replacement at 2’-OH contacts residues. n =3, error bars are S.E.M. (**B**) Ribose replacement at specific residues combined with PS linkages identifies an RNA-free design, FM8.15, with high editing. n =3, error bars are S.E.M. (**C**) Cell-based EGFP editing using wild-type or Y450A and F105A Cas9. n =3, error bars are S.E.M. (**D**) CRISPR activation of an EGFP reporter driven by a TetON promoter using unmodified (crTREa) or modified crRNAs targeting the EGFP promoter. Percent EGFP(+) cells was quantified by flow cytometry. n =3, error bars are S.E.M. (**E)** On-target and predicted off-target editing quantified nanopore amplicon sequencing and using crRNAs bearing spacers targeting endogenous AAVS1, TRAC, and ATR genes. “ON” indicates the intended target while “OFF1/2” are predicted off-targets. n =3, error bars are S.E.M.

To achieve high serum stability, we created and tested a series of crRNAs with full ribose replacement. Although the 2’-F modification performed best in individual walks (**Fig. 3A**), its incorporation at positions 2 and 6 (crEG-FM5) reduced editing to less than 25% of the unmodified crEG control (**Fig. 6B**). We reasoned that introduction of sugar modifications adjacent to these positions might alter R-loop dynamics and potentially improve activity. However, DNA (crEG-FM6), which should increase flexibility, and 2’-OMe (crEG-FM7), which might decrease flexibility, both abrogated editing. In contrast, incorporation of PS linkages, known to reduce hybridization affinity and increase dynamics ^92–95^, flanking positions 1, 2, and 6 (crEG-FM8) restored editing to ∼50% of unmodified crEG levels (**Fig. 6B**). Conversely, increasing the number of PS linkages in the central region of the crRNA (crEG-FM8.1) led to a substantial loss of editing. This implies that site-specific interactions are crucial and excessive amounts of PS are detrimental.

Molecular dynamics simulations suggested that interactions within the Cas9 bridge helix (BH) are important. Electrostatic interactions between the guide RNA’s phosphate backbone and positively charged amino acids (e.g., arginine and lysine) in the BH are reported to regulate both on target and off target editing efficiency ^96^. Based on this rationale, we generated two modified gRNAs: crEG-FM8.2, in which all predicted interacting phosphates were replaced with PS linkages, and crEG-FM8.3, in which non-interacting phosphates were substituted with PS ^46^. Both designs provided almost no editing activity. These results suggested that site-specific interactions between Cas9 and phosphates of the guide RNA at contact residue positions 2 and 6 were important for modulating activity. To identify optimal PS incorporation, we designed crEG-FM8.4 through crEG-FM8.15 (**Fig. 6B**). By systematic shuffling and reduction of PS linkages and DNA content, we arrived at crEG8.15, an entirely ribose-free design. This construct fully restored editing to that of the unmodified crEG control. Furthermore, crEG8.15 demonstrated robust stability in 50% human serum with almost no degradation after 8 h, comparable to the all-2’-F control crEG-SJ16 (**Fig. S7**). The likely role of 2’-OH contacts is to mediate conformational transitions necessary for achieving a catalytic state. PS linkages, when properly placed adjacent to 2’-OH contact positions, modulate Cas9 conformation. The mechanism may involve reduced hybridization affinity or increased dynamics ^92–95^. Alternatively, other mechanisms beyond electrostatics may also be involved. A previous crystal structure of PS-modified DNA bound to annexin A2 revealed that the sulfur atom engages primarily hydrophobic contacts with arginine side chains ^97^.

### Cas9 mutants probe the influence of site-specific ribose replacement

To further probe the higher sensitivity of ribose replacement at 2′-OH contact positions 2 and 6, we generated two Cas9 point mutants. The 2’-OH groups of contact residues 2 and 6 hydrogen bond to the Cas9 peptide backbone near tyrosine 450 (Y450) and phenylalanine 105 (F105), respectively ^46^. Y450A mutations were previously reported to reduce off-target editing via a bipartite seed checkpoint mechanism ^98^ and F105 has been implicated in stabilization of a guide repeat clasp (GRC) checkpoint mechanism ^77^. Thus, we made Y450A and F105A mutant variants of Cas9 and tested them with heavily or fully modified crRNAs (**Fig. 6C**). Y450A and F105A resulted in decreased editing for unmodified crEG. Heavily modified crEG-FM4 and crEG-FM4.4, as well as fully modified crEG-FM8.15, performed similarly to wild-type Cas9 with F105A but all showed reduced editing with Y450A. Editing with crEG-FM8 and crEG-FM8.14 was reduced as expected, with similar editing using F105A. However, Y450A was particularly detrimental when paired with crEG-FM8 and crEG-FM8.15. Together, these results demonstrate that 2’-OH contacts coordinate with Cas9 amino acids to control catalytic activity and suggest that amino acid changes may be uncovered that can further increase the compatibility of Cas9 with chemically modified crRNAs.

### A ribose-free crRNA retains target binding and supports editing at endogenous loci

To assess target engagement of ribose-free chemically modified crRNAs in cells, we utilized our CRISPRa assay. We found that heavily modified crTREa-FM1 and crTREa-FM4.4 designs, as well as fully modified crTREa-FM8.15, activated EGFP robustly, better than the unmodified crTREa control (**Fig. 6D**). In contrast, crTREa-FM8.14 showed reduced EGFP activation. These results mirrored the gene editing activity of each crRNA, supporting restoration of target engagement as a primary mechanism for the FM8.15 design.

To extend our results beyond an EGFP reporter, we synthesized crRNAs with the FM1, FM4.4, FM8.14, and FM8.15 chemical modification schemes targeting endogenous AAVS1, TRAC-tgt (TRAC), and ATR genes. Editing was assessed by nanopore sequencing at on-target and candidate off-target loci (**Fig. 6E**). Editing of AAVS1 and TRAC was reduced for all designs compared to unmodified crRNA. In contrast, all designs except FM8.14 provided editing similar to unmodified crRNA at the ATR gene. The ribose-free FM8.15 design offered about 2/3 and 1/3 the editing efficiency of unmodified crRNA for AAVS1 and TRAC while providing full editing for ATR. Both FM4.4 and FM8.15 provided excellent fidelity, with similar or lower off-target editing compared to unmodified crRNA. Interestingly, the FM1 design increased off target editing in some cases. Together, these results demonstrate that a ribose-free crRNA chemical modification scheme, FM8.15, can provide favorable on- and off-target editing, establishing a promising design to build upon for therapeutic applications.

## Discussion

The 2’-hydroxyl (2’-OH) group is essential for guide RNA (gRNA) function, but its presence prevents full chemical modification for increased stability, posing a significant hurdle for therapeutic development ^70, 99^. The specific regulatory mechanism within the Cas9 protein that necessitated 2’-OH groups remained a puzzle ^70^, leading therapeutic development efforts to focus on alternative delivery strategies rather than full chemical replacement ^53, 100^. In this study, we sought to discover the mechanism underlying a 2’-OH requirement. We focused on Cas9 as a model due to its high degree of mechanistic characterization and its role as a leading platform for CRISPR-based therapeutics. Furthermore, to simplify chemical modification and experimentation, we also focused on crRNA for its small size and predicted site-specific 2’-OH contacts with Cas9. By systematically testing biochemical activity, RNP stability, and target binding and editing in cells, as well as structural investigations with cryo-EM and MD simulations, we found that the 2’-OH contact residues in crRNA are critical for Cas9 target engagement. Specifically, loss of 2’-OH contacts trap Cas9 in conformational states that are not competent for efficient target DNA binding. Cryo-EM showed a reduction in particles that formed the DNA-bound “closed” conformation. To map defects in the conformational and energetic landscape, MD simulations were performed using a modified crRNA with a single DNA substitution that severely compromises binding and catalysis. We conclude that crRNA 2’-OH contacts with the peptide backbone function as critical checkpoints for the allosteric activation of Cas9. Loss of 2’-OH contacts induces a cascade of events, beginning with altered bridge helix (BH) interactions and ending with compromised RuvC and HNH active states.

Chemical modification schemes, including individual position walks with multiple chemistries, demonstrated the importance of 2’-OH contacts between crRNA and Cas9. A general trend that was similar for most chemical modifications, and independent of sequence, supported the hypothesis of a mechanism intrinsic to the Cas9 complex, not to a specific chemistry. However, some chemical ribose replacements were more tolerated than others at individual positions. Based on these chemical probing results and data supporting defects in conformational state transitions, we surmised that full ribose replacement might be achievable if chemistry can be found that enables the requisite conformational changes. Thus, we created a series of fully chemically modified crRNAs, focusing on modifications that might increase or decrease interaction dynamics with Cas9 at or near 2’-OH contact residues. The careful placement of PS linkages at these sites resulted in discovery of the FM8.15 crRNA design, which fully restored editing of EGFP in cells.

The most sensitive contact positions, which we modulated with PS chemistry, interact with the Cas9 backbone near amino acids Y450 and F105. Y450 is a key residue involved in the bipartite seed checkpoint that can reduce off target editing when mutated to alanine ^98^. F105 is an amino acid involved in hydrophobic packing to stabilize another proposed checkpoint that senses RNP assembly and influences target specificity, the guide repeat clasp (GRC) checkpoint ^77^. Mutating both Y450 and F105 to alanine altered editing efficiency when comparing heavily and fully modified crRNAs, supporting a potential overlap of 2’-OH contacts with other checkpoint mechanisms. When tested against three endogenous genes, we observed favorable on- and off-target editing profiles for the FM8.15 design, although efficiency was impacted by target sequence. We propose that the sulfur substitution in the PS linkage modulates dynamics of Cas9 by increasing hybridization dynamics with target DNA and increasing interaction stability with positively charged Cas9 residues, mimicking the role of the 2’-OH and enabling transition to a catalytically competent state.

These findings align with several studies showing that crRNAs lacking 2’-OH groups at critical contacts can in some cases preserve *in vitro* Cas9 cleavage despite a severe loss of cell-based editing ^43, 50, 101^. A reduction in the fraction of Cas9 RNP complexes able to assume a target binding competent state, as observed here, helps explain this phenomenon. When combined with the lower effective RNP concentration inside cells, as well as more challenging chromatin DNA targets, Cas9 complexes that do not efficiently engage target DNA will exhibit reduced gene editing efficiency. *In vitro*, however, slow target association kinetics or fast off rates may be overcome by the high concentration of RNP components and target DNA often used. This perspective clarifies why some crRNAs with ribose replacement chemistries can fail in a cellular context despite performing sufficiently in *in vitro* cleavage assays.

Our findings elucidated the underlying mechanism for the requirement of 2’-OH contacts between crRNA and Cas9, which facilitated the use of combined linkage and sugar chemistries to circumvent the barrier to ribose replacement. This work, however, has some limitations. We focused solely on Cas9 crRNA. The applicability of the combined linkage and sugar chemistry approach to Cas9 tracrRNA or sgRNA, which also require 2′-OH contacts for function ^38, 42–45^, is still unclear. Likewise, how well this approach will translate to other RNA-guided enzymes that require 2’-OH contacts, including Cas12a ^48, 70^, will require more investigation. The mixed performance of the FM8.15 design against endogenous loci demonstrates that it is not yet a universal solution. Developing a broadly applicable method will likely require exploring additional chemical modification combinations in the future, either with PS or more diverse linkages ^102–104^. These may include phosphoramidate, phosphorodithioate, stereo-pure phosphorothioate, methylphosphonate, or mesyl phosphoramidate (MsPA) ^104^. Further studies are also needed to better understand the role of 2’-OH contacts in checkpoint mechanisms, which should include additional cryo-EM, MD simulations, Cas9 mutations, and single molecule studies^85^.

The mechanism by which 2’-OH contacts control Cas9 represents a novel and previously unknown checkpoint for target binding. It signals to Cas9 that the guide RNA is properly poised for target engagement then subsequently mediates the conformational allostery necessary for catalysis. This mechanism may extend to other RNA-guided enzymes and also appears to impact target specificity. These principles provide a clear mechanistic framework for further rational optimization of Cas9, either through Cas9 mutagenesis or guide RNA engineering. The FM8.15 design is a reliable starting point, showing favorable editing and specificity profiles. The stability conferred by complete ribose replacement now makes it possible to perform *in vivo* studies that explore new strategies for initiating and controlling gene editing. Separate delivery of Cas9 and crRNA, which is essential for DNA binding and gene editing, could enable higher safety profiles. For example, better tissue-specific targeting, sequential or multiplexed therapeutic gene editing, and better control over the duration of editing activity. In summary, our findings reveal an unexpected role for 2’-OH contacts in mediating Cas9 target binding and offer a mechanistic underpinning to guide complete ribose replacement for improved therapeutic development of CRISPR-Cas9 and other CRISPR-based drugs.

## Data availability

All data presented is available upon request or can be found in the data repositories described following provided accession numbers.

## Acknowledgements

We thank the many lab members from each contributing research group for helpful discussions. Funding was provided by grants from the National Institutes of Health (7R01GM135646) to K.T.G and M.J.D.

## Author contributions

A.A.P. designed and performed biochemical and cell-based experiments, including gene editing, CRISPRa, *in vitro* cleavage, target DNA binding, and on- and off-target editing and sequencinq, as well as analyzed data, prepared figures, and prepared the manuscript. H.M.B. performed custom crRNA syntheses and purifications. S.S. performed MD simulations and computational modeling and prepared figures and the manuscript. R.C. performed gene editing, CRISPRa, and on- and off-target editing experiments, generated Cas9 mutants, and helped prepare RNP complexes for cryo-EM. S.K.J. performed crRNA syntheses and purifications; M.A.P. created the inducible EGFP Tet-On 3G HeLa cell line used for CRISPRa. M.S.B. performed and analyzed nanopore sequencing for on- and off-target editing. J.A.G.-M. carried out *in vitro* time-course cleavage assays. S.B.E. and C.L.B. assisted with cell-based experiments, while C.A.B. performed additional in vitro assays. L.A., E.M.-A., and D.O. contributed to crRNA synthesis and purification. S.K. conducted cryo-EM data collection and analyses and helped prepare and edit the manuscript. P.I.P. supervised molecular dynamics and computational modeling efforts and prepared figures and the manuscript, M.J.D. supervised crRNA syntheses and prepared and edited the manuscript. K.T.G. conceived the project, guided all experimental design, execution, and interpretation, performed serum stability assays, prepared figures, and wrote and edited the manuscript.

---

**Fig. S1.**
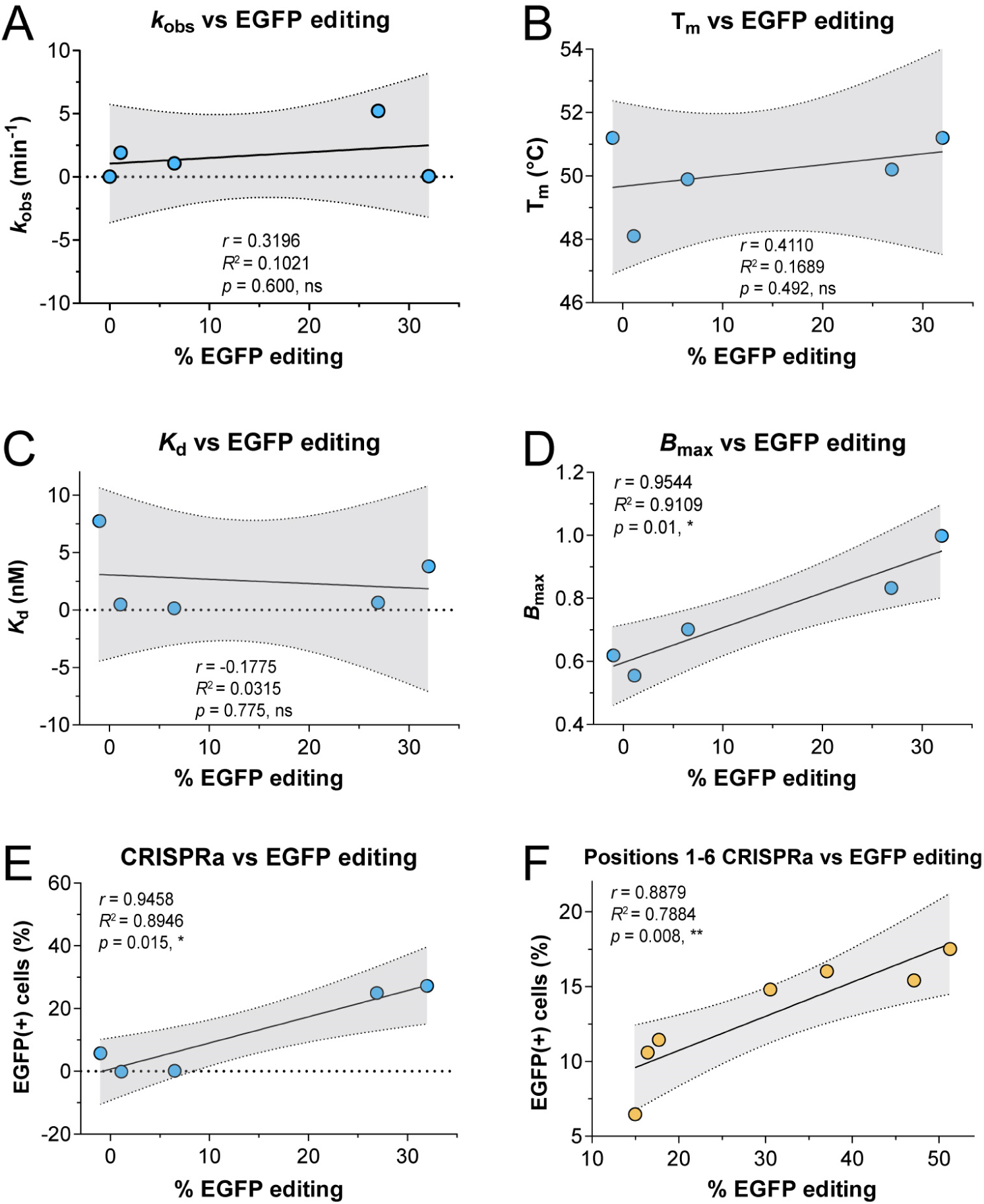
Pearson correlation between EGFP cell editing and other metrics. (**A**) Cleavage kinetics *k*_obs_, (**B**) Melting temperature (*T*_m_), (**C**) Radiolabeled dot blot *K*_d_, (**D**) radiolabeled dot blot *B*_max_, and (**E**, **F**) CRISPRa EGFP(+) cells plotted against % EGFP editing to obtain Pearson’s correlation analyses.

**Fig. S2.**
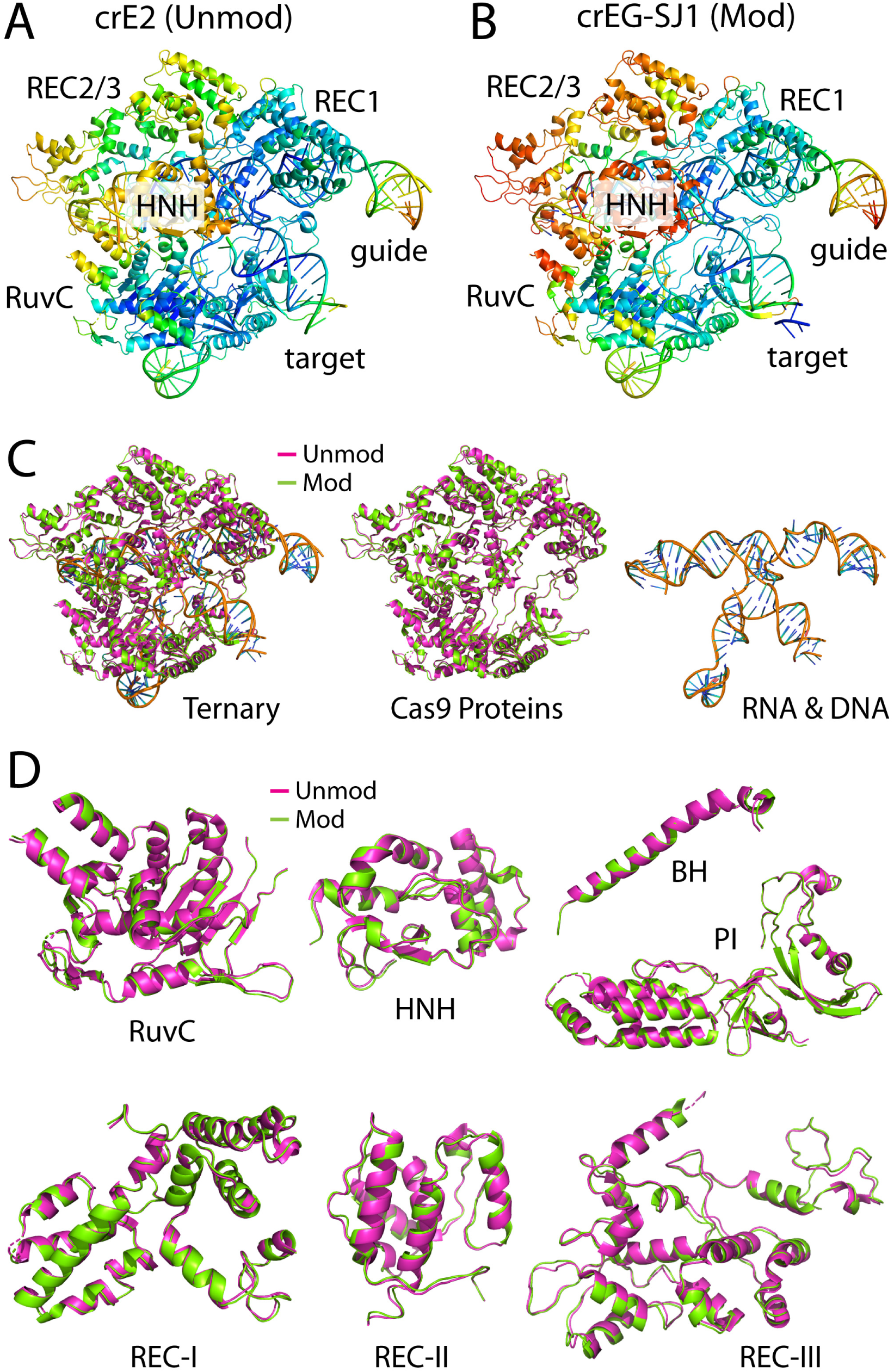
Cryo EM reconstructions and superimpositions of Cas9 ternary complexes with unmodified and chemically crRNAs. (**A**, **B**) Cryo EM reconstructions of SpCas9 ternary complexes assembled with (**A**) unmodified crRNA crE2 or (**B**) 2′ modified crEG-SJ1. Models are colored by atomic B factor from blue (low) to red (high). (**C**) Superposition of the two ternary complexes, with the unmodified (crE2) complex in magenta and the modified (crEG-SJ1) complex in green. Left, full ternary complexes; middle, Cas9 proteins only; right, crRNAs only. (**D**) Close up views of individual Cas9 domains from the superimposed structures, highlighting minimal local perturbations in RuvC, HNH, bridge helix (BH), PI, REC I, REC II and REC III. Unmodified and modified complexes are shown in magenta and green, respectively.

**Fig. S3.**
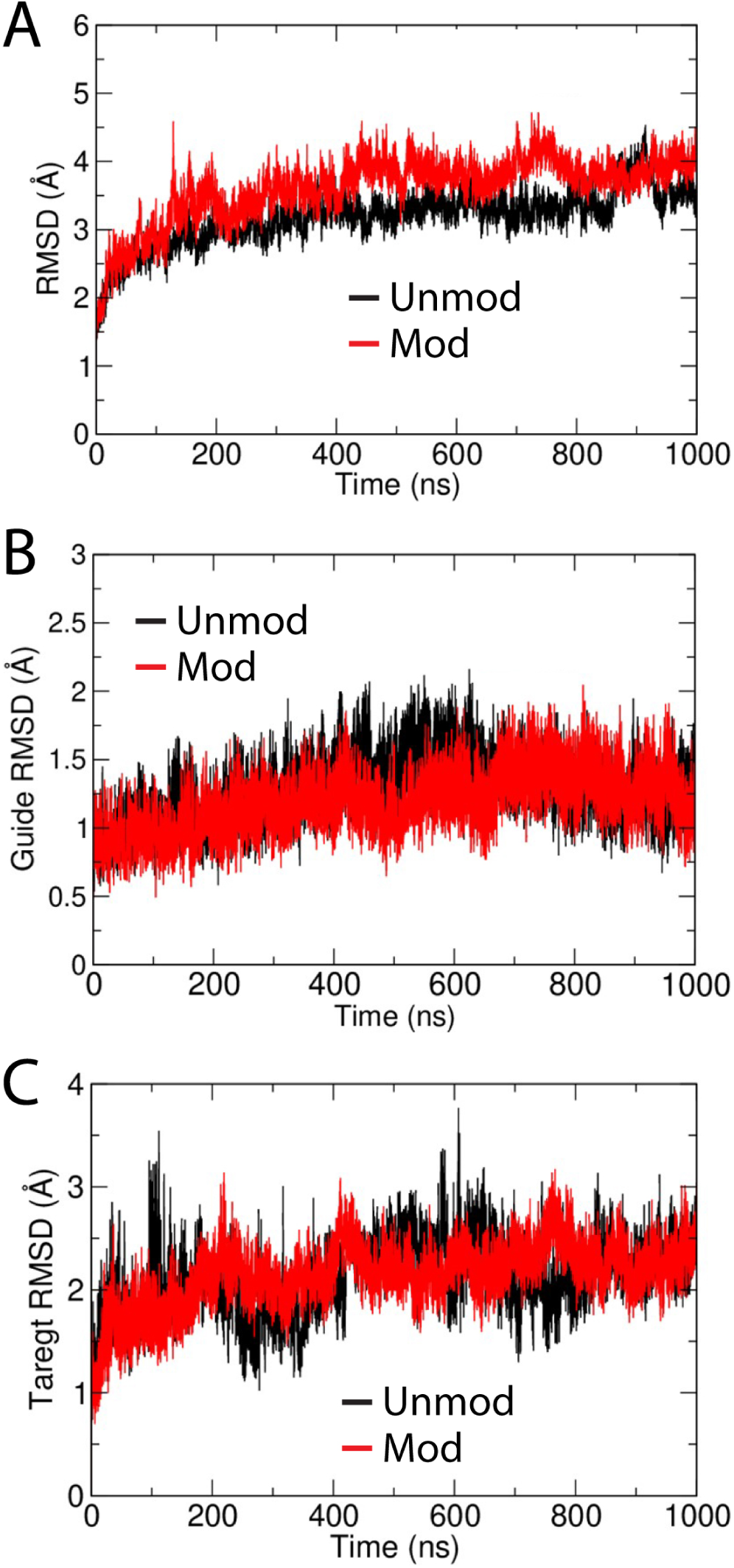
RMSD analysis of Cas9 complexes containing unmodified (crEG) and DNA-modified (crEG_d1) crRNA. Root mean square deviation (RMSD) as a function of simulation time for (**A**) the Cas9 protein backbone, (**B**) the guide RNA backbone, and (**C**) the target DNA backbone. Traces correspond to simulations with unmodified (black) or modified (red) crRNA. RMSD values were calculated from 1 µs molecular dynamics trajectories.

**Fig. S4.**
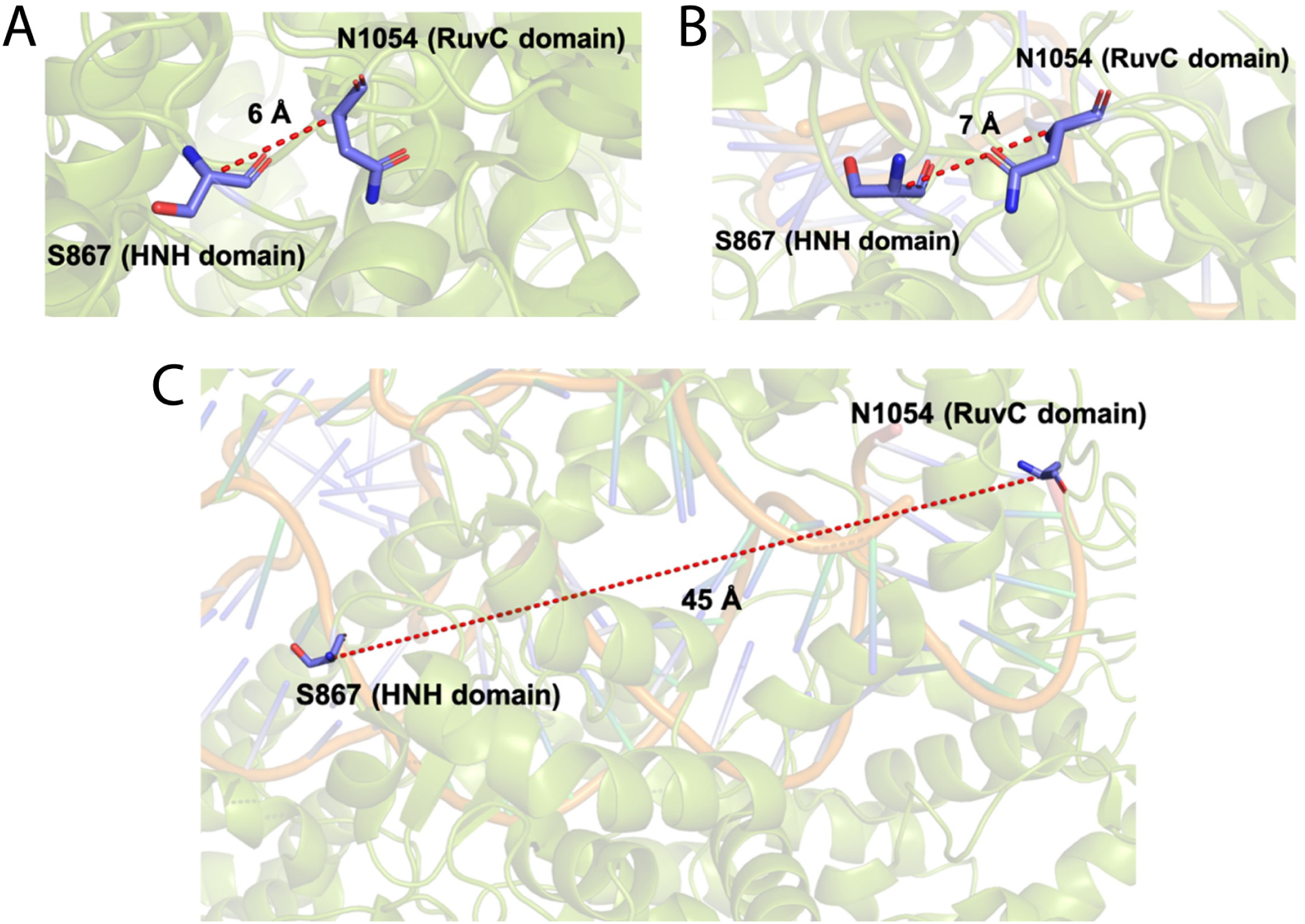
Representative images showing the distance between Cα atoms of S867 and N1054 in the (**A**) Apo (PDB ID: 4CMQ); (**B**) RNA bound form (PDB ID: 4ZT0); and (**C**) DNA bound precatalytic state of Cas9 (PDB ID: 5F9R).

**Fig. S5.**
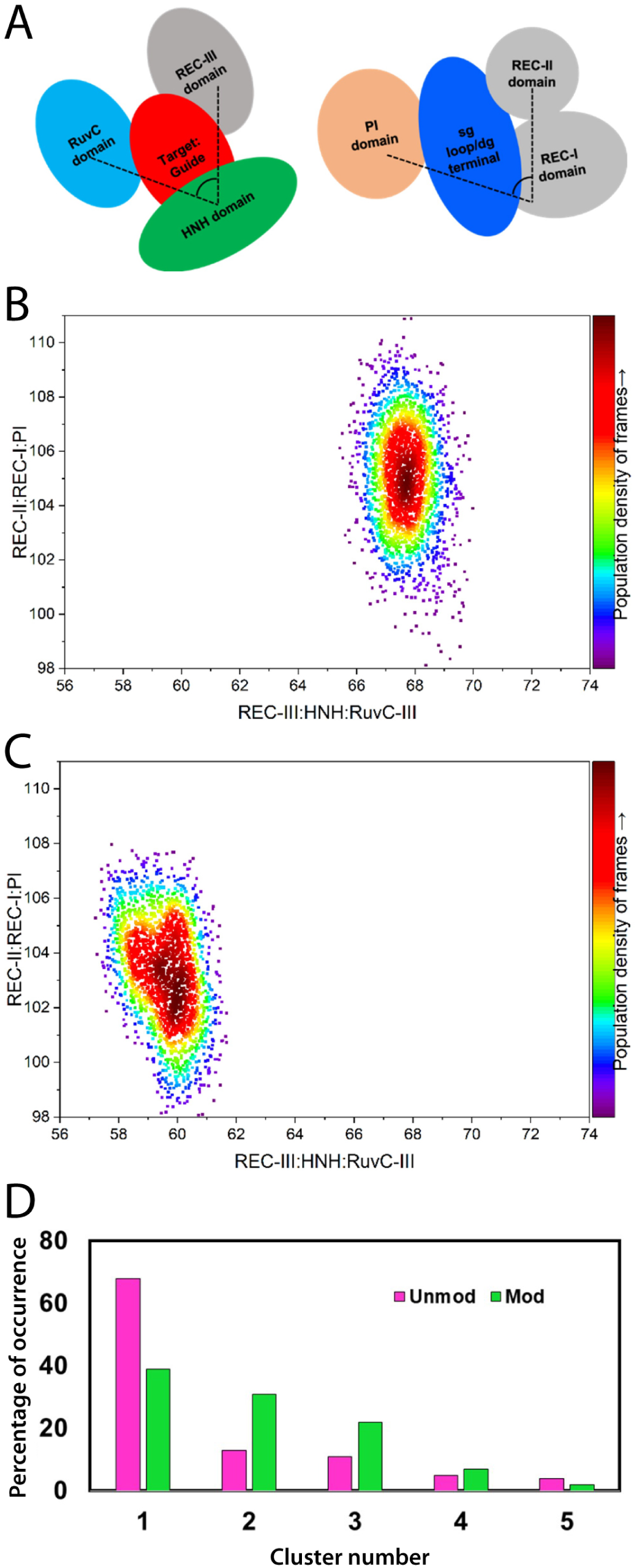
Cas9 interdomain organization and conformational ensembles with unmodified and DNA-modified guide RNA. (**A**) Cartoon representation of Cas9 domain arrangement around the guide and target duplex, indicating the REC-I, REC-II, REC-III, HNH, RuvC-III and PI domains used for interdomain angle measurements. (**B, C**) Two dimensional distributions of interdomain angles between REC-II:REC-I:PI (y-axis) and REC-III:HNH:RuvC-III (x-axis) for complexes containing (**B**) unmodified crEG (Unmod) or (**C**) modified crEG_d1 (Mod). The color scale reports the population density of simulation frames sampled over 1 µs molecular dynamics trajectories. (**D**) Cluster analysis of interdomain angle conformations showing the percentage of frames assigned to each cluster for complexes with unmodified (magenta) or modified (green) crRNA.

**Fig. S6.**
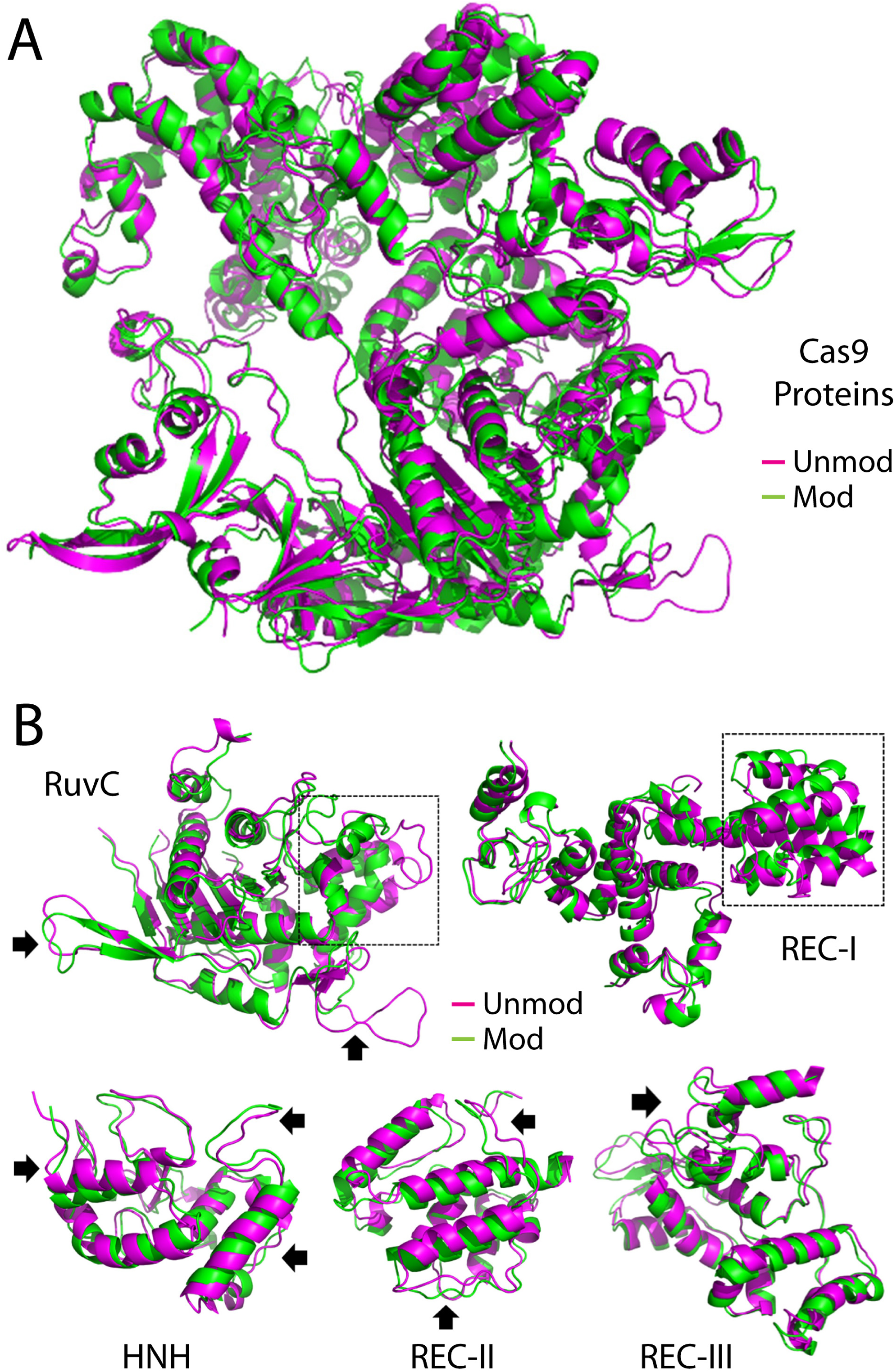
Molecular dynamic structural comparison of dominant Cas9 conformations with unmodified and modified crRNA. (**A**) Superposition of Cas9 structures from the major molecular dynamics clusters for complexes containing unmodified crEG (Unmod) in magenta or modified crEG_d1 (Mod) in green. (**B**) Close up views of superimposed individual domains (RuvC, REC-I, HNH, REC-II and REC-III) from the same clusters, highlighting helices (dashed boxes) and loop (arrows) rearrangements that shift between Unmod and Mod complexes.

**Fig. S7.**
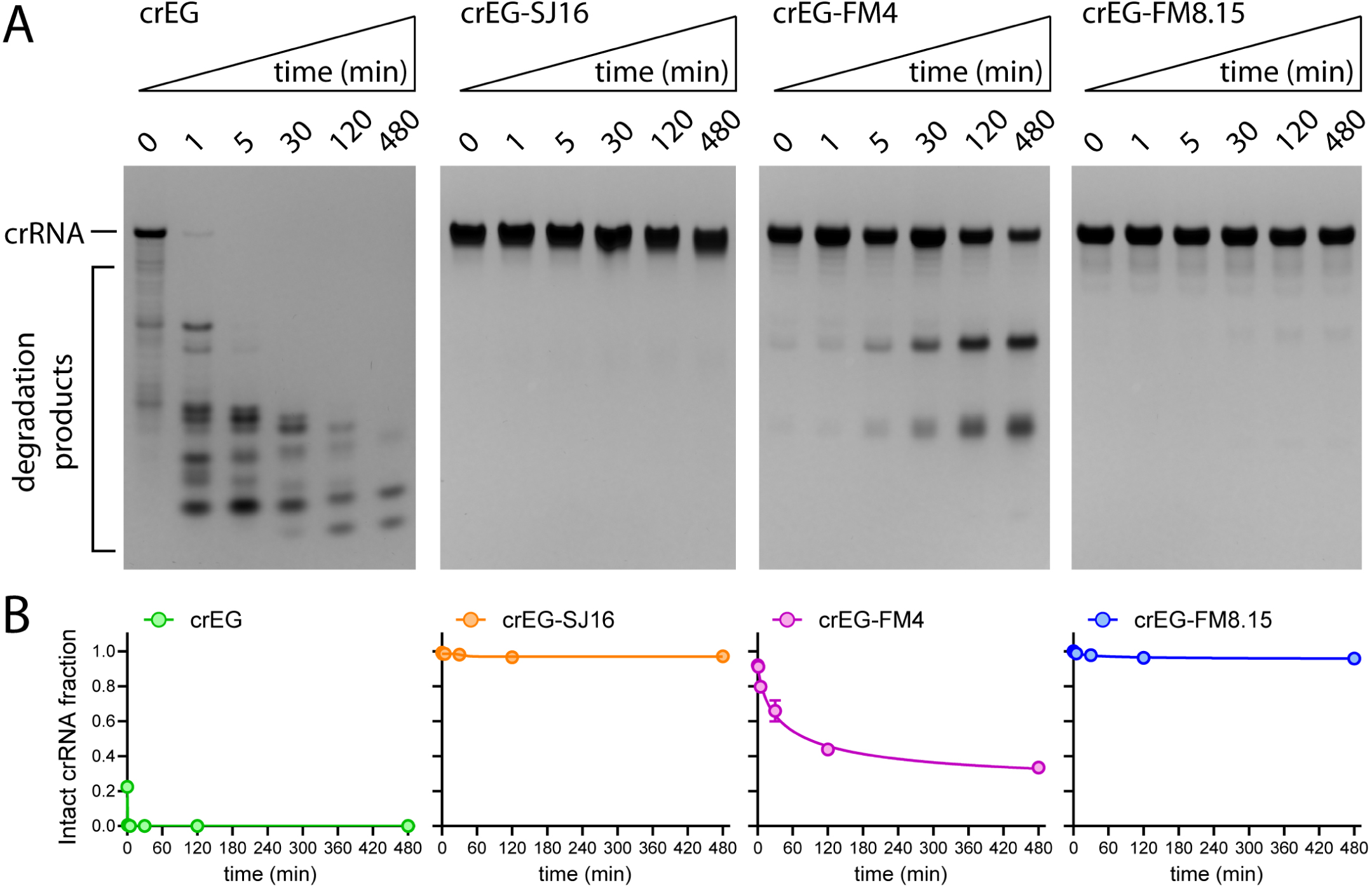
Serum stability assays for select crRNAs. Representative urea-PAGE gel for each crRNA after treatment with 50% human serum for the indicated durations. Quantification of crRNA degradation is shown below each gel. n = 3 experimental replicates, error bars are S.E.M.

